# Differential *EDS1* requirement for cell death activities of plant TIR-domain proteins

**DOI:** 10.1101/2021.11.29.470438

**Authors:** Oliver Johanndrees, Erin L. Baggs, Charles Uhlmann, Federica Locci, Henriette L. Läßle, Katharina Melkonian, Kiara Käufer, Joram A. Dongus, Hirofumi Nakagami, Ksenia V. Krasileva, Jane E. Parker, Dmitry Lapin

## Abstract

Toll/interleukin-1 Receptor (TIR) domains are integral to immune systems across all domains of life. TIRs exist as single-domain and as larger receptor or adaptor proteins. In plants, TIRs constitute N-terminal domains of nucleotide-binding leucine-rich repeat (NLR) immune receptors. Although TIR-NLR and TIR signaling requires the Enhanced disease susceptibility 1 (EDS1) protein family, TIR domains persist in species that have incomplete or no EDS1 members. To assess whether particular TIR groups appear with EDS1, we searched for TIR-EDS1 co-occurrence patterns. Using a large-scale phylogenetic analysis of TIR domains from 39 algae and land plant species, we identify four conserved TIR groups, two of which are TIR-NLRs present in eudicots and two are more widespread. Presence of one TIR-only protein group is highly correlated with EDS1 and members of this group elicit *EDS1*-dependent cell death. By contrast, a more widely represented TIR group of TIR-NB-WD40/TPR (TNP) proteins (formerly called XTNX) has at least one member which can induce *EDS1*-independent cell death. Our data provide a new phylogeny-based plant TIR classification and identify TIR groups that appear to have evolved with and are dependent on *EDS1*, while others have *EDS1*-independent activity.

**One sentence summary:** Land plants have evolved four conserved TIR groups

## Introduction

Toll/interleukin-1 Receptor (TIR) domains regulate immune signaling and cell death in bacteria, animals and plants (Leulier & Lemaitre, 2008; Nimma *et al*., 2017; Bayless & Nishimura, 2020; Ofir *et al*., 2021). In bacteria, TIR domain proteins likely constitute antiphage defense systems or act as virulence factors (Coronas-Serna *et al*., 2020; Morehouse *et al*., 2020; Ofir *et al*., 2021). In animals, TIRs function as signal transduction modules within specialized adaptors (e.g. Myeloid differentiation primary response 88 (MyD88)) and in receptor proteins such as Toll-like receptors (TLRs) and Sterile alpha and Toll/interleukin-1 receptor motif-containing protein 1 (SARM1), which sense pathogen-associated molecular patterns (PAMPs) or cell metabolic changes (O’Neill & Bowie, 2007; Figley *et al*., 2021). In plants, intracellular immune receptors with N-terminal TIR domains have a central domain called nucleotide-binding adaptor shared by APAF-1, certain *R*-gene products, and CED-4 (NBARC or NB-ARC) and C-terminal leucine-rich repeats (LRRs). This receptor class (referred to as TIR-NLR or TNL) detects pathogen virulence factor (effector) activities to induce defenses which often culminate in localized host cell death (Jones *et al*., 2016). A number of plant truncated TIR-only and TIR-NB proteins also contribute to pathogen detection or defense amplification (Nishimura *et al*., 2017; Zhang, X *et al*., 2017b; Bayless & Nishimura, 2020; Tamborski & Krasileva, 2020; Tian *et al*., 2021). No functional TIR adaptors were found in plants and bacteria to date.

Interactions between activated animal TLRs and TIR adaptor proteins transduce pathogen recognition into defense via protein kinase activation and transcriptional reprogramming (O’Neill & Bowie, 2007; Fields *et al*., 2019; Clabbers *et al*., 2021). Importance of these homo- and heterotypic TIR interactions for immune responses is highlighted by the fact that bacterial pathogens in mammals utilize TIR effector hetero-dimerization to disrupt MyD88-mediated TLR signaling (Cirl *et al*., 2008; Yadav *et al*., 2010; Nanson *et al*., 2020). Another TIR mechanism was discovered in human SARM1, in which self-associating TIRs hydrolyze NAD^+^ leading to neuronal cell death (Gerdts *et al*., 2015; Essuman *et al*., 2017; Horsefield *et al*., 2019; Sporny *et al*., 2019). NAD^+^ cleavage activity was also found in TIRs of the bacterial antiphage Thoeris system and TIR-STING cyclic dinucleotide receptors (Morehouse *et al*., 2020; Ofir *et al*., 2021), bacterial TIR effectors (Coronas-Serna *et al*., 2020; Eastman *et al*., 2021), plant TNLs and TIR-only proteins (Horsefield *et al*., 2019; Wan *et al*., 2019; Ma *et al*., 2020). TIR NADase activity and associated host cell death require a conserved catalytic glutamate residue in a pocket formed by the TIR DE interface, BB-loop and the αC-helix in interacting TIRs (Essuman *et al*., 2017; Essuman *et al*., 2018; Horsefield *et al*., 2019; Wan *et al*., 2019; Ma *et al*., 2020; Martin *et al*., 2020; Burdett *et al*., 2021). Thus, TIRs can regulate host-parasite interactions and cell death via NAD^+^ hydrolysis-dependent and independent processes. Many TIR domains are bifunctional enzymes with the capacity for 2’,3’-cAMP/cGMP synthetase activity utilizing RNA and DNA substrates in addition to their NADase activity (Yu *et al*., 2021).

Previously, TIRs in prokaryotes and eukaryotes were divided into 37 groups through Bayesian partitioning with pattern selection (BPPS) (Toshchakov & Neuwald, 2020). The majority of plant TIRs were assigned to three plant-specific groups following domain architectures of the full-length proteins, although ∼1000 plant TIRs remain unclassified (Toshchakov & Neuwald, 2020). The largest plant-specific group was enriched for TIRs from TNLs, and the two remaining groups included TIR-only proteins and TIRs fused to NBARC-like domains (Toshchakov & Neuwald, 2020). The latter group corresponds to so-called XTNX proteins, where X indicates conserved N-terminal and C-terminal sequences (Meyers *et al*., 2002; Nandety *et al*., 2013; Zhang *et al*., 2016). Because XTNXs contain WD40- and tetratricopeptide-like repeats (TPRs) instead of LRRs (Shao *et al*., 2019), we call XTNXs from here on <u>T</u>IR-<u>NBARC</u>-like-β-<u>p</u>ropeller WD40/T<u>P</u>Rs (TNPs), to reflect their domain architecture and fitting with existing NLR nomenclature. The BPPS grouping of plant TIRs aligns with earlier studies employing phylogeny-based group assignment of TIRs (Meyers *et al*., 2002; Nandety *et al*., 2013).

In dicot plants, all tested TIR-only and TNL proteins function via a plant-specific protein family consisting of Enhanced disease susceptibility 1 (EDS1), Phytoalexin-deficient 4 (PAD4) and Senescence-associated gene 101 (SAG101) (Lapin *et al*., 2020; Dongus & Parker, 2021). The EDS1 family proteins contain an N-terminal lipase-like domain and C-terminal α-helical bundle EDS1-PAD4 domain (EP, PFAM: PF18117) with no clear homology outside the EDS1 family (Wagner *et al*., 2013; Baggs *et al*., 2020; Lapin *et al*., 2020). EDS1 forms a dimer with either PAD4 or SAG101 to mediate pathogen resistance and cell death triggered by plant TIRs (Wagner *et al*., 2013; Nishimura *et al*., 2017; Bhandari *et al*., 2019; Gantner *et al*., 2019; Horsefield *et al*., 2019; Lapin *et al*., 2019; Wan *et al*., 2019; Lapin *et al*., 2020; Sun *et al*., 2021). By contrast, expression of the human SARM1 TIR domain or *Pseudomonas syringae* HopAM1 TIR effector in wild tobacco (*Nicotiana benthamiana*; *Nb*) triggered *EDS1*-independent cell death (Horsefield *et al*., 2019; Wan *et al*., 2019; Eastman *et al*., 2021), suggesting a degree of specificity in translating plant TIR catalytic activity into immune responses via the EDS1 family. Consistent with plant EDS1 family – TIR cofunctions, expanded TNL repertoires correlate with the presence of EP domain sequences in seed plants (Wagner *et al*., 2013; Lapin *et al*., 2019; Baggs *et al*., 2020; Liu *et al*., 2021). However, the existence of TNPs and other TIRs in plant genomes that lack *EDS1* (Meyers *et al*., 2002; Gao *et al*., 2018; Toshchakov & Neuwald, 2020) raises the question whether a subset of plant TIRs also function in an *EDS1*-independent manner.

Our aim here is to find signatures of EDS1-TIR co-occurrence which could be used to predict EDS1-dependency of distinct TIR domain groups in plants. By phylogeny-based clustering of predicted TIR sequences from 39 species representing diverse groups of green plants, we identify four TIR groups that are shared by at least two plant lineages. Two of these groups match TIRs of the previously identified TNPs and conserved TIR-only proteins (Meyers *et al*., 2002; Nandety *et al*., 2013). Two other TIR groups belong to TNLs in Angiosperms. *Nb* tobacco mutants for *TNP*s, encoding the most conserved TIR proteins in plants, behaved like wild type in tested PAMP-triggered and TNL immunity outputs and *TNP* genes were unresponsive in the analyzed immune-related expression assays, suggesting immunity-independent *TNP* functions. We further establish that a TNP from maize elicits *EDS1*-independent cell death in *Nicotiana tabacum* transient expression assays. Conversely, immunity-induced expression of the conserved *TIR-only* genes, dependency of cell death elicited by monocot conserved TIR-only proteins on *EDS1* in *Nb*, and co-occurrence with EDS1/PAD4 in angiosperms suggest the importance of an EDS1/PAD4 – conserved TIR-only signaling node in the immune system of flowering plants. Hence, there appears to be selectivity at the level of EDS1 in plant TIR downstream signaling and cell death activity, which fits patterns of TIR domain phylogeny.

## Results

### Land plants have evolved four conserved TIR groups

To study the distribution of TIRs in plants, we utilized predicted protein sequences from 39 species comprising unicellular green algae, non-seed land plants, conifers, and seven clades of flowering plants (*Amborella trichopoda* or *Amborella* hereafter, *Nymphaeales*, *Magnoliids*, *Ceratophyllales*, monocots, superrosids and superasterids) (Supplementary Table 1). In total, 2348 TIRs were predicted using hidden Markov models (HMMs, see Materials and Methods). The number of predicted TIR-containing sequences per plant species ranged from a single protein in *Marchantia polymorpha* (Bowman *et al*., 2017) and *Selaginella moellendorffii* to 435 and 477 in the Rosid *Eucalyptus grandis* and conifer *Pinus taeda*, respectively. Generally, the highest numbers of predicted TIR-containing proteins were found in eudicots (Supplementary Figure 1a; (Sun *et al*., 2014; Liu *et al*., 2021)). Analyses of the protein domain composition revealed 1020 TNLs, 401 TN and 572 TIR-only architectures (Supplementary Figure 1b-d; note TNPs were excluded from these calculations; (Sun *et al*., 2014)). As expected, TNLs were missing in monocots and *Erythranthe guttatus* (Shao *et al*., 2016; Liu *et al*., 2021) and low TNL numbers were found in two Caryophyllales (*Amaranthus hypochondriacus* and *Beta vulgaris*) as inferred previously (Shao *et al*., 2016; Lapin *et al*., 2019; Baggs *et al*., 2020; Liu *et al*., 2021). Whereas TNLs were found in 20 of 39 analyzed species, TIR-only proteins (sequences shorter than 400 amino acids and without other predicted PFAM domains) were present in 33 of 39 species, including unicellular green algae and monocots (Supplementary Figure 1d; (Sun *et al*., 2014; Liu *et al*., 2021)). Thus, TIR-only is likely the most widely adopted TIR protein architecture across plants.

To refine categorization of plant TIRs, we constructed a maximum likelihood (ML) phylogenetic tree for the 2348 TIR sequences (Supplementary Figure 2a, Supplementary Files 1, 2). This analysis revealed four TIR groups shared by several but not all groups of land plants. Algal sequences did not form a monophyletic group and did not fall into the four shared TIR groups. Since algal TIR sequences tended to have long branches, we excluded them from further analysis and repeated the ML tree inference for the remaining 2317 TIR sequences (Supplementary Figure 2b, Supplementary Files 3-5). The same four phylogenetically distinct TIR groups appeared as shared by land plant lineages (Supplementary Figure 2b). A large excess of sequences over number of alignment patterns can lead to false phylogenetic inferences. Therefore, we prepared a reduced ML tree for 307 representative TIRs (Figure 1a) selected from the major groups present on the bigger ML tree (Supplementary Figure 2c, Supplementary Files 6, 7). The same four TIR groups were recovered again, despite different alignments and underlying evolutionary models (Figure 1a; BS>90%, SH-aLRT>80), suggesting that categorization of these four TIR groups is consistent across analyses. Since NBARC domain types correlate with NLR groups (Shao *et al*., 2016; Tamborski & Krasileva, 2020), we tested whether the TIR groups identified here are associated with different NBARC variants. For that, we constructed an ML phylogenetic tree for associated NBARC sequences from full-length TIR-containing sequences used in Figure 1a (Supplementary Figure 3, Supplementary Files 8, 9). NBARCs from sequences within the TIR groups also formed well-supported branches (BS>90%, SH-aLRT>80), suggesting a degree of specificity between conserved TIRs and NBARCs. We conclude that land plants have four phylogenetically distinct TIR groups shared by at least two taxonomic clades.

**Figure 1.**
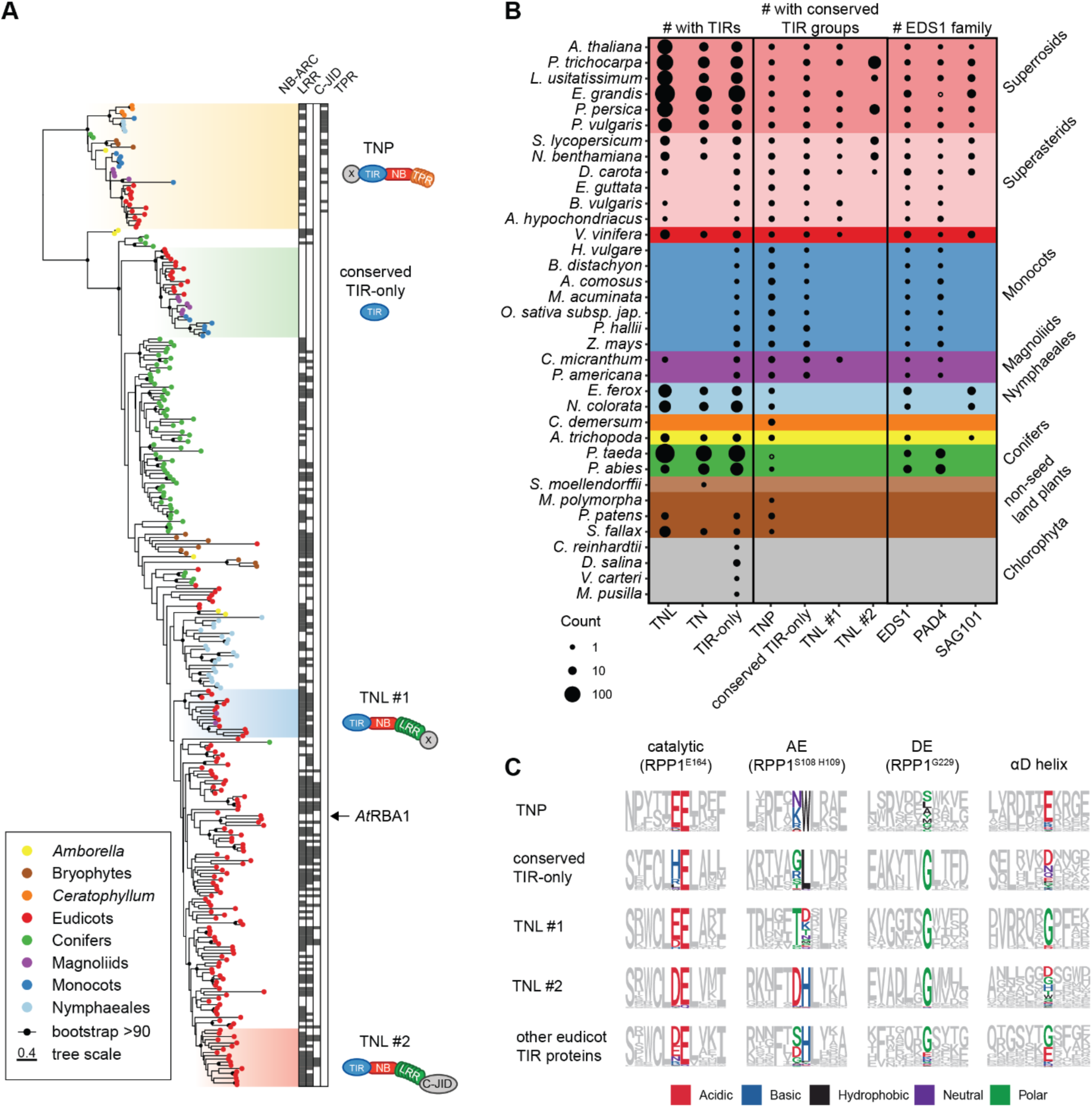
Land plants have four conserved TIR groups. **A** ML tree (evolutionary model WAG+F+R7) of 307 predicted TIR domain sequences representing major TIR families across plant species (full 2317 sequence tree in Supplementary Figure 2b). Branches with BS support ≥90 are marked with black dots. Conserved groups with TIRs from more than one species are highlighted with colored boxes and their predominant domain architecture is depicted. Additional domains predicted in the TIR proteins are annotated as black boxes next to each TIR protein (used HMM listed in Supplementary Table 2). Four most conserved TIR domain groups were named after the predominant domain architecture of their full-length proteins. The TIR-only *At*RBA1/*At*TX1 does not belong to conserved TIR-only proteins. **B** Counts of predicted full-length TIR proteins, proteins with conserved TIRs, and EDS1 family predicted in the species analyzed within this study. TNPs are not included in the counts of TNL, TN and TIR-only proteins. TIR-only proteins are defined as sequences shorter than 400 amino acids, without other predicted PFAM domains. Sizes of circles reflect the counts. *Pinus taeda* has one TNP ortholog that was not identified by HMMs, but via reciprocal BLAST. *Eucalyptus grandis* has a fragment of PAD4-like sequence as determined by TBLASTN searches. **C** Comparison of important TIR domain motifs across the four conserved plant TIR groups. Full sets of TIR domains were taken based on phylogeny (tree in Supplementary Figure 2). Sequence motifs were generated for each TIR group to show conservation of the catalytic glutamate, AE and DE interfaces, as well as residues in the αD helix. *At*RPP1^WsB^ TIR domain was taken as reference. Chemical attributes of the important amino acids are annotated in different colors.

### Conserved TIR groups match full-length sequences with different domain architectures

Next, we investigated whether full-length proteins within each TIR group have specific domain architectures and how these align with earlier studies. Two conserved TIR groups match two TNL families. One of them is also known as “conserved TNL lineage” or “NLR family 31” in studies deploying NBARC phylogeny and synteny searches (Zhang *et al*., 2016; Liu *et al*., 2021). We use the term TNL #1 hereafter for this TNL group. The post-LRR C-terminal extension in TNL #1 proteins does not show similarity to other PFAM domains (Supplementary Figure 2b). Since TNL #1 proteins are found in the majority of eudicots and in the monocot-sister magnoliid *Cinnamomum micranthum* (Zhao *et al*., 2021) but not in conifers, *Amborella* or Nymphaeales (Figure 1b), this TIR group likely emerged in mesangiosperms before the split of monocots and dicots and was likely lost in monocots.

TNLs with the second conserved TIR nested in the NBARC phylogeny-based NLR group called “NLR family 10” in (Zhang *et al*., 2016). We refer to this NLR family 10-nested TNL group as “TNL #2” (Figure 1a). Our TIR phylogenetic grouping did not find evidence for this conserved TIR in *Arabidopsis thaliana* (*Arabidopsis* from here onwards) and *Amborella*. However, reciprocal BLASTP searches with the respective full-length TNL from domesticated tomato (Solyc01g102920.2.1) suggest that these species have one orthologous sequence each (in *Arabidopsis* - AT5G36930). Because we define sequence groups based on TIR conservation, *Arabidopsis* and *Amborella* TNLs do not fall into the TNL #2 group. In contrast to TNL #1 present in 1-4 copies per genome, the TNL #2 group expanded in some eudicot genomes (e.g., 54 genes in poplar) (Figure 1b, S2b; (Zhang *et al*., 2016)). It comprises ∼50% of predicted TNLs in poplar, wild tobacco and tomato. We detected the post-LRR C-terminal jelly-roll/Ig-like domain (C-JID, PFAM: PF20160) in some TNL #2 (Supplementary Figure 2b, Figure 1a; (Van Ghelder & Esmenjaud, 2016; Ma *et al*., 2020; Martin *et al*., 2020; Saucet *et al*., 2021)). Because the C-JID contributes to LRR-specified effector recognition in TNL receptors (Ma *et al*., 2020; Martin *et al*., 2020), we assume that many TNLs in the TNL #2 group have a role in pathogen detection.

The third TIR group (we refer to as conserved TIR-only) corresponds to a small family of ∼200 aa-long proteins with a TIR-only architecture and 1-4 gene copies per genome. This group is present in 22 analyzed magnoliids, monocots, and eudicots but absent in conifers, *Amborella* or *Nymphaeales* (Figure 1b), suggesting its emergence in mesangiosperms similar to the TIR of TNL #1. Strikingly, and in contrast to TNL #1, conserved TIR-only proteins are present in monocots. *Arabidopsis* TX3 and TX9 (Nandety *et al*., 2013) fall into this TIR group. We noticed that the TIR-only effector sensor protein Recognition of HopBA1 (RBA1) does not belong to this conserved TIR-only group (Figure 1a; (Nishimura *et al*., 2017)). Therefore, we conclude that the TIR-protein domain architecture is not a suitable basis for assigning TIR types.

The most taxonomically widespread plant TIR-containing proteins are TNPs ((Figure 1b); (Meyers *et al*., 2002; Zhang, Y-M *et al*., 2017)). TNPs are almost ubiquitous in analyzed species including the aquatic flowering plant duckweed *Wolffia australiana* with a reduced NLR repertoire (Figure 1b, Supplementary Figure 4, Supplementary Files 10, 11; (Zhang, Y-M *et al*., 2017; Baggs *et al*., 2020; Michael *et al*., 2020; Liu *et al*., 2021)). The TNP group matches *Arabidopsis* TN17-like and TN21-like sequences (Nandety *et al*., 2013). Structure-guided comparison with NBARCs from characterized plant NLRs revealed NBARC-characteristic motifs Walker A, RNBS-B, Walker B, RNBS-C, GLPL and MHD, but not the RNSB-B-located TTR/TTE motif (Ma *et al*., 2020; Martin *et al*., 2020) in TNP NBARCs (Supplementary Figure 5). We confirmed that the TIR and NBARC-like sequence in TNPs is followed by C-terminal TPRs that are sometimes picked up WD40 HMMs (Figure 1a; (Meyers *et al*., 2002; Nandety *et al*., 2013; Zhang, Y-M *et al*., 2017; Shao *et al*., 2019)). Since a TIR-NBARC-WD40 architecture is present in red algae *Chondrus crispus* (Gao *et al*., 2018), we tested orthology of TIRs in this algal group to green plant TNPs. Both reciprocal BLAST searches and grouping based on the TIR ML phylogenetic tree (Supplementary Figure 6, Supplementary Files 12, 13) suggest that TIRs from *C. crispus* TIR-NBARC-WD40 sequences are not orthologous to TNP TIRs.

Taken together, the four conserved TIR groups matched predicted full-length proteins with different architectures, but these domain architectures are insufficient to define individual TIR groups.

### Glutamate in the NADase catalytic motif is shared by four conserved TIR groups

We assessed whether key residues critical for plant TIR functions are present in the four conserved TIR groups, utilizing primary sequence and secondary structure-informed alignments. The SH motif is central to an AE dimerization interface in the TIR domains of TNL Resistant to *Pseudomonas syringae* 4 (RPS4) (Williams *et al*., 2014; Zhang, X *et al*., 2017a). This motif did not show a high level of sequence conservation across the four conserved TIR types (Figure 1c). A glycine residue that is necessary for TIR self-association via a second DE interface and for cell death and NADase activity of *Brachypodium distachyon Bd*TIR and *Arabidopsis* RBA1 TIR-only proteins (Nishimura *et al*., 2017; Zhang, X *et al*., 2017a; Wan *et al*., 2019) was conserved in all tested TIR groups except the TNPs (Figure 1c). AlphaFold2 structures of conserved TIR-only proteins from rice and *Arabidopsis* are predicted to differ from known plant TIRs in a poorly structured region in place of the TNL TIR-characteristic αD-helices (see rice TIR-only - grey, *Arabidopsis* – purple in Supplementary Figure 7) (Bernoux *et al*., 2011). This αD-helical region is important for cell death activities of TNL receptors RPS4 (Sohn *et al*., 2014) and L6 (Bernoux *et al*., 2011) and for 2′,3′-cAMP/cGMP synthetase activity found in several plant TIR domains (Yu *et al*., 2021). The glutamate residue which is indispensable for TIR NADase activity (Essuman *et al*., 2018; Horsefield *et al*., 2019; Wan *et al*., 2019; Ma *et al*., 2020) was present in all four conserved TIRs (Figure 1c), pointing towards a possible NAD^+^ hydrolytic activity of these TIR groups.

### Conserved TIR groups show different co-occurrence patterns with EDS1 family members

Since the EDS1 family connects plant TIR activity to resistance and cell death outputs in dicot plants (Lapin *et al*., 2020), we tested whether the distributions of EDS1 family members and identified conserved TIR groups align across species. We therefore built an ML tree for 200 sequences with an EP domain that uniquely defines the EDS1 family (Supplementary Figure 8, Supplementary files 14 and 15, PFAM PF18117) and inferred numbers of EDS1, PAD4 and SAG101 orthologs per species (Supplementary Table 3, Figure 1b). As expected, EDS1 and PAD4 were present in most seed plants while SAG101 was not detected in conifers, monocots and Caryophyllales (Figure 1b, Supplementary Figure 8, (Lapin *et al*., 2019; Baggs *et al*., 2020; Liu *et al*., 2021). Of the four TIR groups, the conserved TIR-only type showed highest correlation with EDS1 and PAD4 in mesangiosperms (Figure 1b), indicating a possible functionally conserved TIR-only-EDS1/PAD4 signaling module. By contrast, TNPs were present in non-seed land plants and aquatic plants that do not have the *EDS1* family genes (Figure 1b and S4; (Baggs *et al*., 2020)), pointing to EDS1-independence of TNP activities.

The above co-occurrence analyses confirmed that the TNL #1 group has a SAG101-independent distribution in angiosperms (Liu *et al*., 2021). This prompted us to search for other protein family orthogroups (OGs) that co-occur with TNL #1 and SAG101 (Supplementary Figure 9). Using Orthofinder, we built OGs for predicted protein sequences from ten species. Five species (*Oryza sativa, Ananas comosus, Picea abies*, *Erythranthe guttata, Aquilegia coerulea*) lacked SAG101 and TNL #1 (Figure 1b, (Zhang *et al*., 2016; Liu *et al*., 2021)). One species (*A. hypochondriacus*) had TNL #1 but no SAG101. Finally, we included four species (*A. thaliana, E. grandis, Populus trichocarpa, Solanum lycopersicum*) with SAG101 and TNL #1. We imposed a strict co-occurrence pattern to retain only high confidence candidates. Seven and five OGs followed the SAG101 and TNL #1 distribution, respectively. These findings were refined using reciprocal BLAST searches in genomes of the discriminatory species *Beta vulgaris* (*TNL#1^+^/SAG101^-^;* (Lapin *et al*., 2019; Liu *et al*., 2021))*, Sesamum indicum* and *Striga hermonthica* (*TNL#1^-^*/*SAG101*^-^; (Shao *et al*., 2016; Liu *et al*., 2021)). After this filter, two OGs showed co-occurrence with SAG101 - *Arabidopsis* hypothetical protein AT5G15190 and arabinogalactan proteins AT2G23130/AT4G37450 (AGP17/AGP18) (Supplementary Figure 9). The other two OGs that co-occurred with the conserved angiosperm TNL #1 had *Arabidopsis* terpene synthase 4 (TES, AT1G61120) and glutaredoxins ROXY16/17 (AT1G03020/AT3G62930) as representatives (Supplementary Figure 9). The functions of these genes in TIR-dependent defense are unknown. Overall, we concluded that conserved TIR groups show different distribution patterns in flowering plants and their co-occurrence with SAG101 is limited.

### Conserved *TIR-only* genes are transcriptionally induced in immune-triggered tissues

The broad species distributions of the four plant TIR groups prompted us to investigate their patterns of gene expression across species. Public RNAseq data for seven plant species including *Arabidopsis thaliana*, *Nicotiana benthamiana*, *Hordeum vulgare,* and *Marchantia polymorpha* were used (Figure 2a, Supplementary Figure 10). The samples originated from infected or immunity-triggered tissues as well as mock-treated or untreated control samples. Relative transcript abundance of *TNP* genes was generally unresponsive to the treatments in dicots and *Marchantia*, but it was elevated in several monocot expression datasets. *TNLs* from the TNL #1 and TNL #2 groups were induced in pathogen-infected *Nb* samples (Supplementary Figure 10). Most strikingly, the conserved *TIR-only* genes were either not detected or expressed at a very low level in non-stimulated tissues but displayed induction in multiple immunity-triggered samples in both monocot and dicot species (Figure 2a, Supplementary Figure 10). To explore further defense-related expression control of *TIR*-*only* genes, we analyzed time series RNAseq data for *Arabidopsis* with activated bacterial PAMP- or effector-triggered immune signaling (PTI and ETI; Figure 2b, (Saile *et al*., 2020)). Infiltration of the PTI-eliciting non-pathogenic *Pseudomonas fluorescens Pf*0-1 weakly induced the conserved *Arabidopsis TIR-only* gene *AtTX3*. Higher levels of *AtTX3* expression were detected in samples with *Pf*0-1 delivering effectors recognized by NLRs (Figure 2b, (Saile *et al*., 2020)). Taken together, these observations suggest that expression of the conserved *TIR-onl*y genes is responsive to immunity triggers.

**Figure 2.**
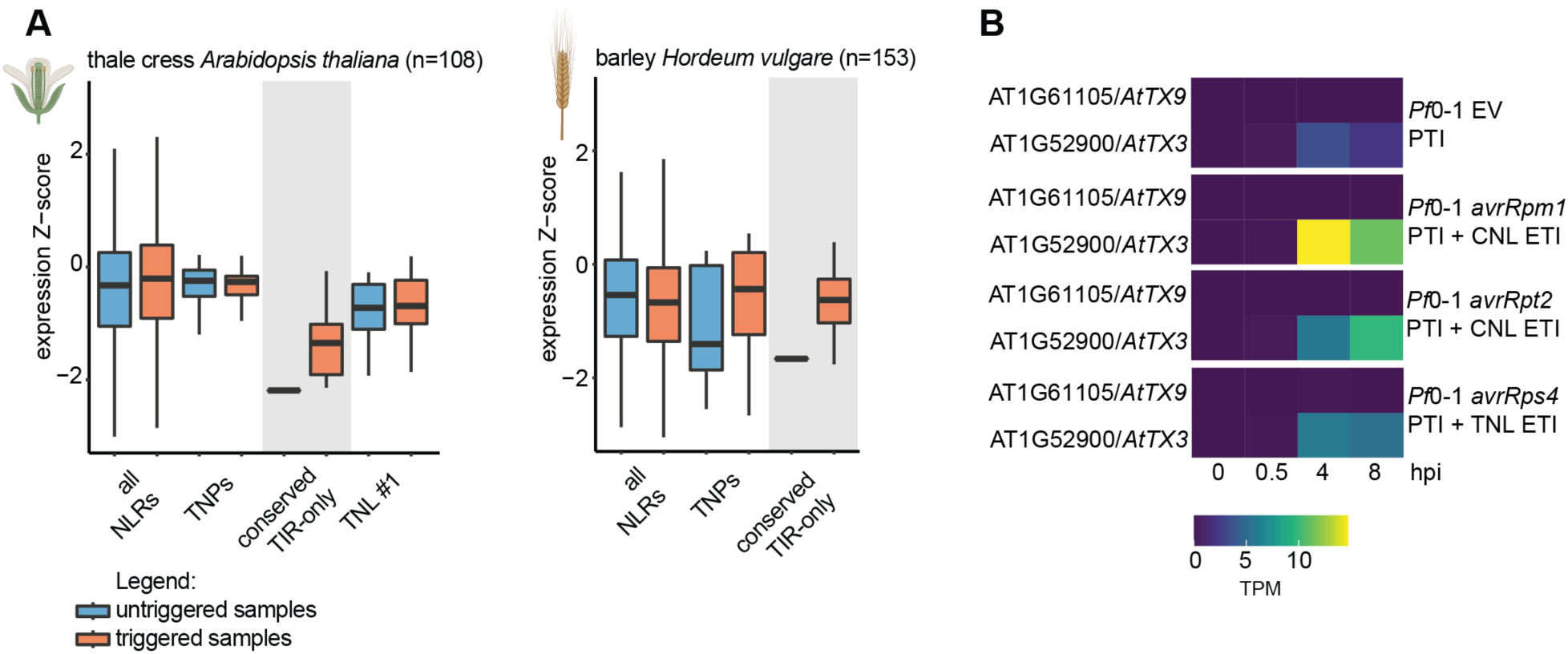
Expression of conserved *TIR-only* genes is upregulated during immune signaling. **A** Comparison of untriggered and immune-triggered expression of *NLRs* and genes corresponding to conserved *TIR* groups in *Arabidopsis thaliana* and *Hordeum vulgare*. Data were taken from publicly available RNAseq experiments including immune-triggered and infected samples. Created with elements from BioRender.com. **B** Heatmaps showing expression of conserved *TIR-only* genes in PTI and ETI trigger combinations in *Arabidopsis*. Expression data were taken from (Saile *et al*., 2020). Triggers include *Pseudomonas fluorescens Pf*0-1 empty vector (EV) for PTI trigger, *Pf*0-1 *avrRpm1*, *Pf*0-1 *avrRpm1* for PTI + CNL ETI and *Pf*0-1 *avrRps4* for PTI + TNL trigger. TPM = transcript per million.

### Monocot conserved TIR-only induce *EDS1*-dependent cell death in *N. benthamiana*

Since the conserved TIR-only proteins co-occur with EDS1 and PAD4 (Figure 1b), we investigated if they trigger *EDS1*-dependent cell death similar to *B. distachyon* conserved TIR-only (*Bd*TIR) (Wan *et al*., 2019). For this, we cloned monocot *TIR*-only genes from rice (*Os*TIR, Os07G0566800) and barley (*Hv*TIR, HORVU2Hr1G039670) and expressed them as C-terminal mYFP fusions in *Nb* leaves using *Agrobacterium*-mediated transient expression assays (Figure 3a). Co-expression of TNL Recognition of *Peronospora parasitica* 1 (RPP1^WsB^) with its matching effector ATR1^Emoy2^ as a positive control (Krasileva *et al*., 2010; Ma *et al*., 2020) resulted in cell death visible as leaf tissue collapse at 3 days post infiltration (dpi) (Figure 3a). mYFP as a negative control did not produce visible cell death symptoms (Figure 3a). Leaf areas expressing rice and barley conserved TIR-only proteins collapsed in *Nb* wild type (WT) at 3 dpi but not in *eds1a* mutant plants (Figure 3a). As the tested monocot TIR-only proteins accumulated in *Nb eds1a* (Figure 3b), we concluded that members of this TIR-only group induce *EDS1*-dependent cell death (Wan *et al*., 2019). The cell death response was fully suppressed in TIR-only mutant variants in which the NADase catalytic glutamate residue was substituted by alanine (*Os*TIR^E133A^ and *Hv*TIR^E128A^; Figure 3a). Similarly, mutation of a conserved glycine at the DE TIR interface which is important for TIR NADase activity (Horsefield *et al*., 2019; Wan *et al*., 2019; Ma *et al*., 2020) fully (*Os*TIR^G188R^) or partially (*Hv*TIR^G183R^), eliminated the cell death response (Figure 3a). All tested TIR-only variants accumulated in *Nb* leaves (Figure 3b). These data show that monocot-derived conserved TIR-only proteins induce host cell death dependent on an intact NADase catalytic site, DE interface and *EDS1* signaling.

**Figure 3.**
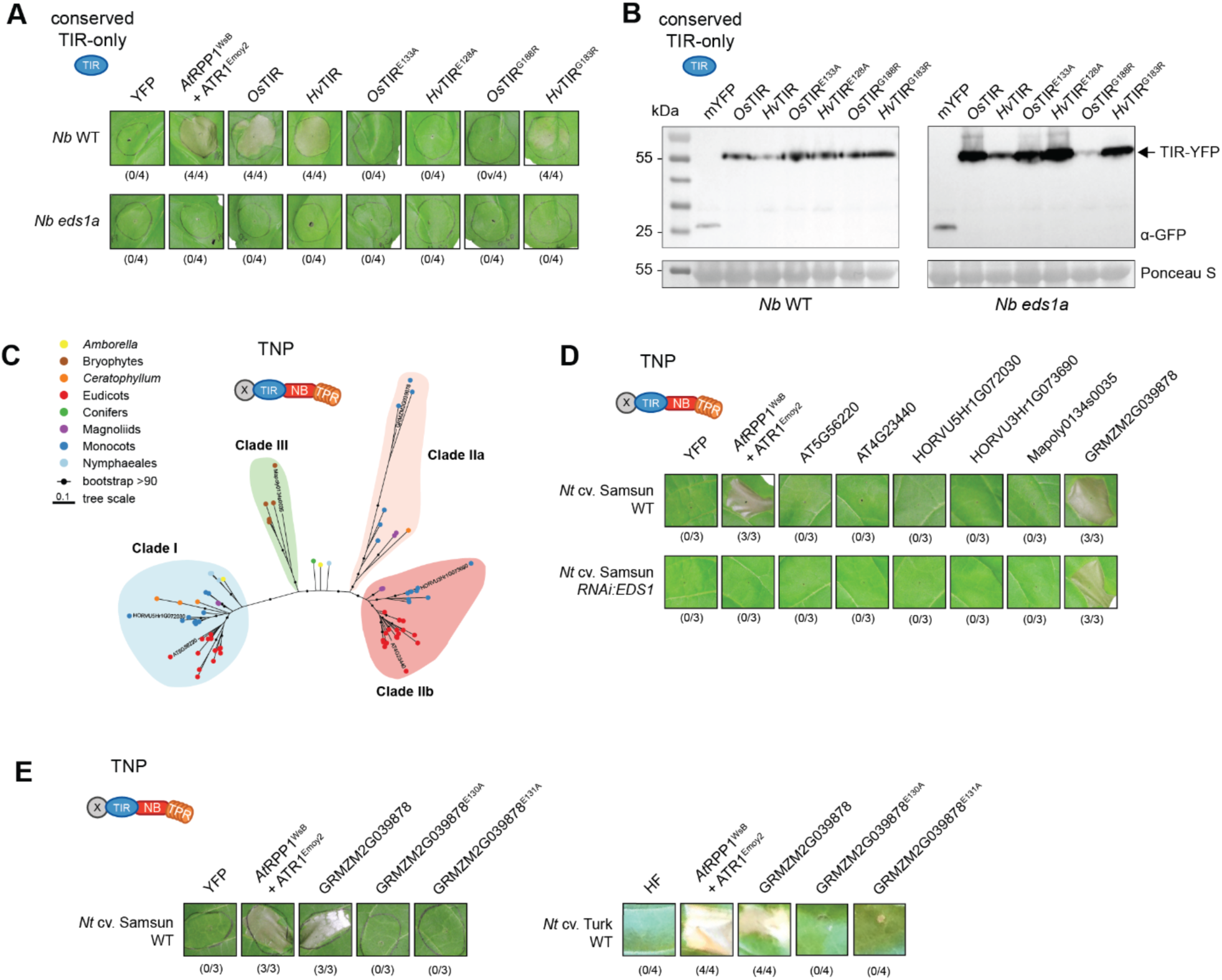
A maize TNP induces *EDS1*-independent cell death in *N. tabacum*. **A** Macroscopic cell death symptoms induced by *Agrobacterium*-mediated overexpression of conserved monocot YFP-tagged TIR-only proteins in *Nicotiana benthamiana* (*Nb*) wild type (WT) and the *eds1a* mutant. Pictures were taken three days after agroinfiltrations. Numbers below panels indicate necrotic / total infiltrated spots observed in three independent experiments. **B** TIR-only protein accumulation in infiltrated leaves shown in **A** was tested via Western Blot. Expected sizes for YFP-tagged TIR-onlys and free YFP as control are indicated. Ponceau S staining of the membrane served as loading control. **C** ML tree (from IQ-TREE, evolutionary model JTT+G4) of 77 predicted TNP NBARC (Supplementary File 18, E<0.01) domains representing the plant species analyzed withing this study. Branches with BS support ≥90 are marked with black dots. The three conserved TNP clades are highlighted with colored boxes. Clade nomenclature was partly adapted from Zhang *et al*. 2017. **D** Macroscopic cell death symptoms induced by *Agrobacterium*-mediated overexpression of different TNP proteins from four major clades (*Arabidopsis thaliana*, *Hordeum vulgare* and *Marchantia polymorpha* TNPs were YFP-tagged, *Zm*TNP HF-tagged) representing members of clades shown in **C** in *Nicotiana tabacum* (*Nt*) c.v. “Samsun” wild type (WT) and the *RNAi:EDS1* knock-down mutant. Pictures were taken five days after agroinfiltrations. Numbers below panels indicate necrotic / total infiltrated spots observed in three independent experiments. **E** Overexpression of *Zm*TNP WT and mutant variants in the two adjacent putative glutamates (E130, E131) in *Nt* cv. “Samsun” and “Turk” WT. Pictures was taken five days after agroinfiltration and repeated three times with similar results. Numbers below panels indicate necrotic / total infiltrated spots observed in three independent experiments.

### A maize Clade IIa TNP induces *EDS1*-independent cell death in *N. tabacum*

TNPs are present in plants regardless of presence of the EDS1 family (Figure 1b, Supplementary Figure 2, Supplementary Figure 4, (Nandety *et al*., 2013; Zhang, Y-M *et al*., 2017)). We therefore hypothesized that TNPs act *EDS1*-independently. On the ML tree for TNP NBARC-like sequences selected with help of a custom built HMM (Supplementary Files 16-18), three major TNP clades were recovered, with one splitting into two smaller subclades (Figure 3c). Clade I, Clade IIa and Clade IIb match previously described TNP clades (Zhang, Y-M *et al*., 2017). Expectedly, Clade IIa is missing from eudicots (Figure 3c, (Zhang, Y-M *et al*., 2017)). All bryophyte TNP sequences formed a separate third clade (Clade III, Figure 3c). We selected representative sequences from the above three TNP clades to test whether they induce cell death: *Arabidopsi*s AT5G56220 and barley HORVU5Hr1G072030 from Clade I, maize GRMZM2G039878 from Clade IIa*, Arabidopsis* AT4G23440 and barley HORVU3Hr1G073690 from Clade IIb, and *Marchantia* Mapoly0134s0035 from the bryophyte-specific Clade III (Figure 3c). The C-terminally tagged (maize TNP was 6xHis-3xFLAG (HF)-tagged, others mYFP-tagged) TNPs were expressed in leaves of tobacco *Nicotiana tabacum* (*Nt*) cv. Samsun or a corresponding RNAi:*EDS1* line (Duxbury *et al*., 2020) using *Agrobacterium*-mediated transient expression assays. We scored cell death visually as collapse of the infiltrated area at 5 dpi, using co-expression of *At*RPP1^WsB^-mYFP with effector ATR1^Emoy2^ as a positive control for *EDS1*-dependent cell death (Figure 3d). Expression of maize Clade IIa *Zm*TNP (GRMZM2G039878), but not other TNP forms, consistently resulted in cell death which was *EDS1*-independent (Figure 3d). We were unable to detect any of the TNP proteins on immunoblots. To test whether the predicted maize TNP NADase catalytic glutamate, which is conserved across plant TIRs (Figure 1c), is required for cell death, we substituted adjacent glutamate residues E130 or E131 in *Zm*TNP with alanines (*Zm*TNP^E130A^ and *Zm*TNP^E131A^; Figure 3e). Cell death was abolished for both mutant variants in cv. “Samsun” and “Turk” tobacco *Nt* cultivars. We concluded that *Zm*TNP likely induces *EDS1*-independent cell death via its TIR putative catalytic motif, although we cannot exclude that *Zm*TNP^E130A^ and *Zm*TNP^E131A^ proteins are less stable than cell death-promoting wild-type *Zm*TNP.

### *Botrytis*-infected *N. benthamiana tnp* mutants develop smaller necrotic lesions

To explore possible TNP functions, we developed two independent CRISPR-Cas9 single and quadruple *tnp* mutants, respectively, in *Marchantia polymorpha* and *Nb* (Supplementary Figure 11). The *TNP*-less plants displayed a similar morphology to wild-type (Figure 4a and 4b). Hence, despite the high conservation and wide distribution in land plants, *TNP* genes do not appear to be essential for growth of tested land plants under laboratory conditions.

**Figure 4.**
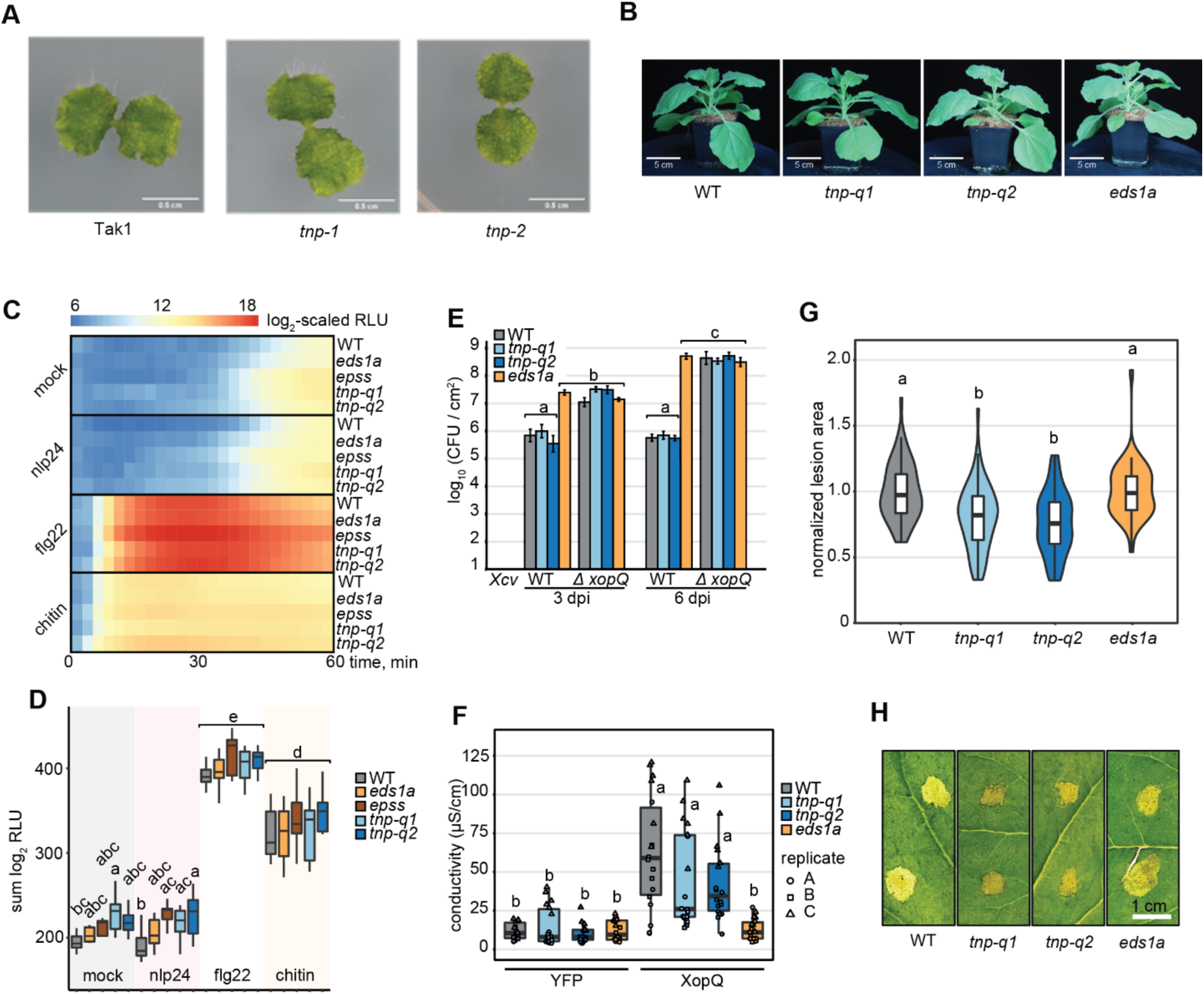
TNPs are not crucial for plant survival but negatively regulate resistance against *Botrytis cinerea* in *N. benthamiana*. **A** Macroscopic images of 2-week-old *Marchantia polymorpha* Tak1 WT and two independent *tnp* CRISPR knockout lines. Genomic sequences of the two *tnp* lines are depicted in Supplementary Figure 11. **B** Side-view images of 4-week-old *N. benthamiana* WT, two independent *tnp* quadruple CRISPR knockout lines (*tnp-q1*, *tnp-q2*) and the *eds1a* mutant. Plants were grown in long-day (16 h light) conditions. Genomic sequences of the two *tnp* quadruple lines are depicted in Supplementary Figure 11. **C** ROS burst upon several PAMP triggers in *N. benthamiana* WT, *eds1a*, *eds1a pad4 sag101a sag101b* (*epss*) and *tnp* quadruple mutants (*tnp-q1*, *tnp-q2*). Values are means of log_2_-transformed RLU after addition of 2 μM nlp24, 200 nM flg22 or 4 mg/ml chitin and were recorded for 60 min, n = 10-12, from three independent biological replicates. **D** Total ROS produced after 60 min PAMP treatment. Values are sums of log_2_-transformed RLU in **C**. Genotype-treatment combinations sharing letters above boxplots do not show statistically significant differences (Tukey HSD, α = 0.05, n = 10-12, from three independent biological replicates). **E** *Xanthomonas campestris* pv. *vesicatoria* (*Xcv*) growth assay in *N. benthamiana*. Plants were syringe-infiltrated with *Xcv* 85-10 (WT) and XopQ-knockout strains (*Δ xopQ*) at OD_600_ = 0.0005. Bacterial titers were determined at three and six days post infiltration (dpi). Genotype-treatment combinations sharing letters above boxplots do not show statistically significant differences (Tukey HSD, α = 0.01, n = 12, from three independent biological replicates). **F** Electrolyte leakage assay as a measure of XopQ-triggered cell death in *N. benthamiana* three days after *Agrobacterium* infiltration (OD_600_ = 0.2) to express XopQ-Myc. YFP overexpression was used as negative control. Genotype-treatment combinations sharing letters above boxplots do not show statistically significant differences (Tukey HSD, α = 0.01, n = 18, from three independent biological replicates). **G** Lesion area induced by *Botrytis cinerea* strain B05.10 infection in *N. benthamiana*. Plants were drop-inoculated with spore suspension (5*10^5^ spores/ml) and lesion areas were measured 48 hours after inoculation. Values shown are lesion areas normalized to WT. Genotypes sharing letters above boxplots do not show statistically significant differences (Tukey HSD, α = 0.01, n = 10-12, from five independent biological replicates). **H** Macroscopic images of *B. cinerea* induced lesions measured in **G**.

Since PTI and TNL ETI readouts are well established for *Nb*, we used the two independent *Nb tnp* CRISPR mutant lines to assess whether *TNP* genes influence defense signaling. Reactive oxygen species (ROS) bursts triggered by PAMPs flg22 or chitin were not altered in the *tnp* mutants (Figure 4c,d) indicating that *TNPs* are dispensable for the PAMP perception and induction of immediate downstream ROS burst. Also, the *Nb tnp* mutants allowed wild type (WT)-like growth of virulent *Xanthomonas campestris* pv. *Vesicatoria* (*Xcv*) bacteria without a XopQ effector that would otherwise trigger TNL Recognition of XopQ (Roq1) (*Xcv ΔXopQ* Figure 4e). In TNL Roq1 bacterial growth assays, the *tnp* mutants were also indistinguishable from resistant WT plants, although the *eds1a* mutant was fully susceptible to *Xcv* (Figure 4e, (Adlung *et al*., 2016; Schultink *et al*., 2017)). Similarly, Roq1 induced cell death was unaffected in the *tnp* mutants after *Agrobacterium*-mediated transient expression of XopQ (Figure 4f), whereas *eds1a* displayed low electrolyte leakage (Figure 4f). Therefore, *TNP*s are dispensable for the tested PTI and ETI outputs in *N. benthamiana*.

We analyzed responses of the *Nb tnp* mutants to infection by the necrotrophic fungus *Botrytis cinerea*. Both *tnp* lines developed smaller necrotic lesions 48 h after spore application while the *eds1a* mutant behaved like WT (Figure 4g,h). The phenotypes of WT and *eds1a* compared to *tnp* mutants when challenged with *Botrytis cinerea* suggest that *Nb TNP*s, directly or indirectly, contribute to *B. cinerea* lesion development via an *EDS1*-independent route.

## Discussion

TIR signaling domains mediate cell death and immune responses across kingdoms, including plants. Here, we analyzed plant TIR conservation and distribution using newly available genomes from major lineages of land plants and the ML phylogenetic tools (Nguyen *et al*., 2015; Chernomor *et al*., 2016) allowing large scale evolutionary analyses. We recovered four interspecific plant TIR groups which so far have no described functions in defense signaling. While two interspecific TIR groups matched TIR-only and TNPs (Meyers *et al*., 2002; Nandety *et al*., 2013; Zhang, Y-M *et al*., 2017; Toshchakov & Neuwald, 2020), two additional TIR groups corresponded to separate angiosperm TNL families. Consistent with differing patterns of co-occurrence with the EDS1 family, conserved TIR-only proteins from monocots and a maize TNP triggered cell death in tobacco, respectively, dependently and independently of *EDS1*. Thus, variation exists in the *EDS1* dependency of plant TIR-promoted cell death.

Although TNL NBARCs of land plants are nested within NBARCs of charophytes (Gao *et al*., 2018), none of the four conserved TIR groups included sequences from unicellular chlorophyte algae (Supplementary Figure 2), red algae *Chondrus crispus* and charophyte *Klebsormidium nitens* (Supplementary Figure 6). Also, our reciprocal BLAST searches did not find TNP orthologs in charophytes *K. nitens* and *Chara braunii*. Hence, the conserved TIR groups of land plants likely did not originate in algae. Specifically, our study suggests that TIR groups from conserved TIR-only and TNL groups #1 and #2 have likely emerged in flowering plants. Major TIR groups of animals (TIRs of TLRs, Myd88, SARM1) are present in *Drosophila melanogaster* and *Homo sapiens* separated by ∼700 million years (confidence interval 643-850 MY) (O’Neill & Bowie, 2007; Toshchakov & Neuwald, 2020). However, a lack of detectable orthology for three of four TIR groups across non-seed land plants (∼500 MY, confidence interval 465-533 MY) (Kumar *et al*., 2017) suggests that plant and animal TIRs may be evolving under different constraints and selection pressures.

We show that the full-length protein domain architecture is insufficient to define TIR groups. Conserved TIR-only proteins form a group that is phylogenetically distinct from TIR-only RBA1 (also known as *At*TX1) and *At*TX12 (Nandety *et al*., 2013; Nishimura *et al*., 2017) which are closer to the TIRs of TNLs RPS4 and Lazarus 5 (LAZ5) (Supplementary Figure 2). *EDS1*-dependent cell death activity of conserved and RBA1/TX-like TIR-only proteins (Figure 3a; (Nishimura *et al*., 2017; Horsefield *et al*., 2019; Wan *et al*., 2019; Duxbury *et al*., 2020)), as well as their immunity-related transcriptional induction (Figure 2; (Nandety *et al*., 2013)), suggest functional convergence of TIR-only groups in plant immunity. Since TIR-only is the most widespread TIR protein architecture in green plants (Supplementary Figure 1; (Sun *et al*., 2014)), structure-informed comparative analyses of different TIR-only groups will be crucial to understand plant immunity networks.

We found differences in copy number of the different TIR group members, with over 50 eudicot TNLs #2 present in the poplar genome in contrast to 1-4 gene copies for the other TIR groups. NLRs have among the highest copy number variation in plants (Baggs *et al*., 2017), ranging from 3,400 NLRs in *Triticum aestivum* (Steuernagel *et al*., 2020) to 1 in *Wolffia australiana* (Michael et al., 2020). High variability in copy number is often associated with the generation of diversity towards a sensor role (Nozawa & Nei, 2008; Kanduri *et al*., 2013; Prigozhin & Krasileva, 2021), whereas conserved low copy number is a feature of signaling components. Presence of the effector-sensing C-JID domain in multiple eudicot TNLs #2 (Figure 1a, Supplementary Figure 2) further suggests they act as pathogen-sensors. It will be interesting to test if conserved TIR-only proteins act as plant TIR adaptor proteins similar to Myd88 and Myd88 adaptor-like cofunctioning with PAMP-recognizing TLRs (O’Neill & Bowie, 2007; Nanson *et al*., 2020).

The absence of conserved TNLs and TIR-only clades in multiple plant species (Figure 1b) suggests that these TIR protein families are not essential for plant viability. TNPs are almost ubiquitous to land plants (Zhang, Y-M *et al*., 2017) and we generated CRISPR-Cas9 mutants of all *TNPs* in *M. polymorpha* and *Nb*. Tobacco quadruple *tnp* mutants and the effectively *TIR*-less *M. polymorpha tnp* mutant were viable and had no obvious growth defects under laboratory conditions (Figure 4). Thus, TNPs and other TIR-containing proteins are likely not essential for plant growth in contrast to Toll and TLR signaling in animals (Anthoney *et al*., 2018), however further research is needed to clarify this.

We found that conserved TIR-only from monocots and a *Zm*TNP triggered cell death in tobacco leaves (Figure 3a,c,d) and this likely required a glutamic acid residue in their conserved catalytic motifs (Figure 1c), as observed in plant TIRs (Horsefield *et al*., 2019; Wan *et al*., 2019). Notably, expression of *Zm*TNP in *N. tabacum* resulted in an *EDS1*-independent cell death, resembling SARM1 (Horsefield *et al*., 2019) and HopAM1 (Eastman *et al*., 2021). These findings suggest that plant TIRs have differing enzymatic activities (Horsefield *et al*., 2019; Wan *et al*., 2019; Duxbury *et al*., 2020) selectively promoting EDS1 family signaling. Given that several plant TIRs were reported to have2′,3′-cAMP/cGMP synthetase activity besides being NADases (Yu *et al*., 2021), it will be interesting to examine the range of enzymatic functions across plant TIR groups and how these affect plant immunity.

## Materials and methods

### Prediction, alignment and phylogenetic analysis of TIRs and other domains

Proteomes of 39 plant species (Supplementary Table 1) were screened for TIR domains using hmmsearch (HMMER 3.1b2, --incE 0.01) with TIR and TIR-related HMMs from the Pfam database (Supplementary Table 2). Redundant TIR sequences found with different TIR and TIR-like HMMs (overlap >20 aa) were removed. Proteins predicted to contain TIR domains were used to build a multiple sequence alignment (MSA) using the MAFFT algorithm (v7.407, fftns or ginsi, with up to 1000 iterations) (Katoh *et al*., 2002). The MSA was filtered and columns with more than 40% gaps were removed using the Wasabi MSA browser (http://was.bi/). The remaining sequences were used to build the maximum likelihood phylogenetic trees with IQ-TREE (version 1.6.12, options: -nt AUTO -ntmax 5 - alrt 1000 -bb 1000 -bnni, (Nguyen *et al*., 2015; Chernomor *et al*., 2016)). The resulting trees were visualized and annotated using the online phylogenetic tree manager iTOL v5 (Letunic & Bork, 2021) or the R package ggtree (Yu, 2020). Sequence data were processed in R with the Biostrings (https://bioconductor.org/packages/Biostrings). Prediction of other domains was performed with hmmsearch (HMMER 3.1b2, --E 0.01) on Pfam A from release 34.0.

### Presence and absence analysis of proteins consistent with SAG101 and conserved angiosperm TNL #1

Orthofinder (v.2.3.11) was run on the following proteomes: P.abies 1.0, Osativa 323 v7.0, Acomosus 321 v3, Acoerulea 322 v3, Ahypochondriacus 459 v2.1, Slycopersicum 514 ITAG3.2, Mguttatus 256 v2.0, Athaliana 167 TAIR10, Egrandis 297 v2.0, Ptrichocarpa 533 v4.1. The *P.abies* proteome was downloaded from congenie.org, all other proteomes were downloaded as the latest version of primary transcript from the Phytozome database (v12) on March 31 2020. Then, we extracted orthogroups that followed the pattern of presence and absence of interest using the following custom scripts extract_orthogroup_TNL_absent_v2.py and extract_orthogroup_SAG101_absent_v2.py. Scripts and orthofinder output are available on github (https://github.com/krasileva-group/TIR-1_signal_pathway.git). *A. thaliana* genes from each orthogroup were searched using tBLASTn against *S. indicum* (Ensembl Plants), *S. hermonthica* (COGE) and *B. vulgaris* (Ensembl Plants). The top hit was then searched with BLASTX or BLASTP (if a gene model was available) back against *A. thaliana* proteome.

### Generation of expression vectors

*TNP* coding sequences without Stop codons were amplified from cDNA (*Arabidopsis thaliana* Col-0, *Hordeum vulgare* cv. Golden Promise, *Oryza sativa* cv. Kitaake, *Marchantia polymorpha* Tak1) using oligos for TOPO or BP cloning (Supplementary Table 4). Coding sequences were amplified with Phusion (NEB) or PrimeStar HS (Takara Bio) polymerases and cloned into pENTR/D-TOPO (Thermo Fisher Scientific) or pDONR221 vectors and verified by Sanger sequencing. Mutations in the sequences were introduced by side-directed mutagenesis (Supplementary Table 4). Recombination of sequences into pXCSG-GW-mYFP (Witte *et al*., 2004) expression vector was performed using LR Clonase II enzyme mix (Life Technologies). *ZmTNP* was synthesized by TWIST Bioscience with codon optimization for expression in *Nicotiana benthamiana;* two fragments were required to synthesize *ZmTNP*. The two fragments were ligated during golden gate cloning into pICSL22011 (oligos listed in Supplementary Table 4) using BsaI restriction sites. Vectors were verified by Sanger sequencing. Site directed mutagenesis of *ZmTNP* was carried out using Agilent technologies QuickChange Lightning Site-Directed Mutagenesis Kit (210518) (oligos listed in Supplementary Table 4). Expression vectors harbouring *At*RPP1^WsB^ and ATR1^Emoy2^ were previously published (Ma *et al*., 2020).

### Transient protein expression and cell death assays in tobacco species

*Agrobacterium tumefaciens* strains GV3101 pMP90RK or pMP90 carrying desired plasmids were infiltrated into *Nicotiana benthamiana or Nicotiana tabacum* leaves at a final OD_600_ of 0.5. For *N. benthamiana* infiltrations, *A. tumefaciens* strain C58C1 pCH32 expressing the viral DNA silencing repressor P90 was added (OD_600_ = 0.1). Prior to infiltration using a needle-less syringe, *A. tumefaciens* strains were incubated in induction buffer (10 mM MES pH 5.6, 10 mM MgCl_2_, 150 nM Acetosyringone) for 1 to 2 h in the dark at room temperature. Protein samples were collected at 2 dpi for Western Blot assays. Macroscopic cell death was recorded using a camera at 3 dpi. For electrolyte leakage assays, six 8 mm leaf disks were harvested for infiltrated leaf parts at 3 dpi and washed in double-distilled water for 30 min. After washing, leaf disks were transferred into 24-well plates, each well filled with 1 ml ddH_2_O. Conductivity of the water was then measured using a Horiba Twin ModelB-173 conductometer at 0 and 6 hours.

### Western blot analysis

To test protein accumulation after *A. tumefaciens* infiltrations in tobacco plants, three 8 mm leaf disks were harvested per protein combination at 2 dpi and ground in liquid nitrogen. Ground tissue was dissolved in 8 M Urea buffer, vortexed for 10 min at RT and centrifuged at 16,000 xg for 10 min (Ma *et al*., 2020). Total protein extracts were resolved on a 10 % SDS-PAGE gel and subsequently transferred onto a nitrocellulose membrane using the wet transfer method. Tagged proteins were detected using primary antibodies (list antibodies) in a 1:5000 dilution (1x TBST-T, 2 % milk (w/v), 0.01 % (w/v) NaAz), followed by incubation with HRP-conjugated secondary antibodies. Signal was detected by incubation of the membrane with Clarity and Clarity Max substrates (BioRad) using a ChemiDoc (BioRad). Membranes were stained with Ponceau S as loading control.

### ROS burst assays in *N. benthamiana*

A ROS burst in response to PAMP elicitors was measured according to (Bisceglia *et al*., 2015). Four-mm leaf discs from 4^th^ or 5^th^ leaves of 5-week-old *Nb* plants were washed in double-distilled (mQ) water for 2h and incubated in 200 μl of mQ water in 96-well plates (Greiner Bio-One, #655075) under aluminum foil overnight. The mQ was then substituted by a solution of L-012 (Merck SML2236, final 180 μM) and horseradish peroxidase (Merck, P8125-5KU, 0.125 units per reaction). Elicitors flg22 (Genscript, RP19986, final 0.2 μM), chitin (from shrimp shells, Merck C7170, resuspended in mQ for 2h and passed through 22 μm filter, final 4 mg/ml), and nlp24 (Genscript, synthesized peptide from *Hyaloperonospora arabidopsidis* NLP3 AIMYAWYFPKDSPMLLMGHRHDWE, crude peptide, final 2 μM) were each added to a 250 μl reaction. Luminescence was recorded on a Glomax instrument (Promega) at 2.5 min intervals. Log_2_-transformed relative luminescence units were integrated across time points for the statistical analysis (ANOVA, Tukey’s HSD test).

### *Xcv* infection assays in *N. benthamiana*

*Xanthomonas campestris* pv. *vesicatoria* (*Xcv*) bacteria was infiltrated in four weeks old *A. N. benthamiana* mutant leaves at a final OD_600_ of 0.0005. The *Xcv* strain carrying XopQ (WT) and one strain lacking XopQ (Δ *xopQ*) were dissolved in 10 mM MgCl_2_. Bacterial solutions were infiltrated using a needleless syringe. After infiltration, plants were placed in a long-day chamber (16 h light/ 8 h dark at 25°C/23°C). Three 8 mm leaf disks representing technical replicates were collected 0, 3 and 6 dpi to isolate the bacteria and incubated in 1 ml 10 mM MgCl_2_ supplemented with 0.01 % Silvet for 1h at 28 °C at 600 rpm shaking. Dilutions were plated on NYGA plates containing 100 mg/L rifampicin and 150 mg/L streptomycin.

### Botrytis infection assays in N. benthamiana

*Botrytis cinerea* strain B05.10 was grown on potato glucose agar (PGA) medium for 20 days before spore collection. Leaves from 4/5-week-old soil-grown *Nicotiana benthamiana* were drop inoculated by placing 10 μl of a suspension of 5 × 10^5^ conidiospores ml^−1^ in potato glucose broth (PGB) medium on each side of the middle vein (4/6 drops per leaf). Infected plants were placed in trays at room temperature in the dark. High humidity was maintained by covering the trays with a plastic lid after pouring a thin layer of warm water. Under these experimental conditions, most inoculations resulted in rapidly expanding water-soaked necrotic lesions of comparable diameter. Lesion areas were measured 48 hours post infection by using ImageJ software.

### Generation of *M. polymorpha tnp* CRISPR/Cas9 mutants

Guide RNA design was performed using CRISPR-P 2.0 (http://crispr.hzau.edu.cn/CRISPR2/) where the sequence of Mapoly0134s0035 was inputted (guide RNAs are listed in Supplementary Table 4). *Marchantia polymorpha* Tak-1 was transformed as described in (Kubota *et al*., 2013) with the exception that *Agrobacterium tumefaciens* strain GV3101 pMP90 was employed. Briefly, apical parts of thalli grown on 1/2 Gamborgs B5 medium for 14 days under continuous light were removed using a sterile scalpel and the basal part of each thallus was sliced in 4 parts of equal size. These fragments were then transferred to 1/2 Gamborgs B5 containing 1% sucrose under continuous light for 3 days to induce calli formation before co-culture with *A. tumefaciens*. On the day of co-culture, *A. tumefaciens* grown for 2 days in 5 ml liquid LB with appropriate antibiotics at 28°C and 250 rpm were inoculated in 5 ml liquid M51C containing 100 µM acetosyringone at an estimated OD_600_ of 0.3-0.5 for 2.5 to 6 hours in the same conditions. The regenerated thalli were transferred to sterile flasks containing 45 ml liquid M51C and *A. tumefaciens* was added at a final OD_600_ of 0.02 in a final volume of 50 ml of medium with 100 µM acetosyringone. After 3 days of co-culture agitated at 400 rpm under continuous light, the thalli fragments were washed 5 times with sterile water and then incubated 30 min at RT in sterile water containing 1 mg/ml cefotaxime to kill bacteria. Finally, plants were transferred to 1/2 Gamborgs B5 containing 100 µg/ml hygromycin and 1 mg/ml cefotaxime and grown under continuous light for 2 to 4 weeks. Successful mutagenesis was validated by PCR amplification (oligos listed in Supplementary Table 4) and subsequent Sanger sequencing. Two independent lines were selected for further experiments.

### Generation of *N. benthamiana tnp* CRISPR/Cas9 mutants

Guide RNA design was performed using CRISPR-P 2.0 (http://crispr.hzau.edu.cn/CRISPR2/) where the four *NbTNP* sequences were inputted (guide RNAs are listed in Supplementary Table 4). *N. benthamiana* WT plants were transformed according to (Ordon *et al*. 2019, dx.doi.org/10.17504/protocols.io.sbaeaie). Successful mutagenesis was validated by PCR amplification (oligos listed in Supplementary Table 4) and subsequent Sanger sequencing. Two homozygous quadruple mutants were selected.

### Analysis of publicly available immune-related RNAseq datasets

RNAseq data (Supplementary Table 5) were downloaded from Sequence Read Archive with sra toolkit (SRA Toolkit Development Team, https://github.com/ncbi/sra-tools; v.2.10.0). After FastQC quality controls (Andrews, S. 2010; A Quality Control Tool for High Throughput Sequence Data; http://www.bioinformatics.babraham.ac.uk/projects/fastqc/), reads were trimmed with Trimmomatic (v0.38, LEADING:5 TRAILING:5 SLIDINGWINDOW:4:15 MAXINFO:50:0.8 MINLEN:36) (Bolger *et al*., 2014). Transcript abundance was quantified with Salmon (v.1.4.0, --fldMean=150 --fldSD=20 for single-end reads, --validateMappings –gcBias for paired-end reads) (Patro *et al*., 2017). The tximport library (v 1.22.0) was used to get the gene expression level in transcript-per-million (tpm) units (Soneson *et al*., 2015). Since RNAseq samples are coming from diverse studies that use different library preparation methods and sequencing platforms, tpm values were standardized per sample and the derived z-scores were used for visualization of the expression levels. Genome versions used as a reference for transcript quantification: *Arabidopsis thaliana* - TAIR10, *Oryza sativa* group *japonica* - IRGSP-1.0, *Hordeum vulgare* - IBSCv2, *Zea mays* - B73v4, *Marchantia polymorpha* v3.1, *Nicotiana benthamiana* v1.0.1. NLR genes were predicted with NLRannotator (https://github.com/steuernb/NLR-Annotator; (Steuernagel *et al*., 2020)).

## Supplementary Data

Supplementary Table 1. List of species used in this study

Supplementary Table 2. List of HMMs used in this study

Supplementary Table 3. Counts of EDS1 family members across species

Supplementary Table 4. Oligonucleotides used in this study

Supplementary Table 5. List of RNAseq accessions

Supplementary File 1. Alignment used to produce ML tree in Supplementary Figure 2a

Supplementary File 2. ML tree in Supplementary Figure 2a (Newick format)

Supplementary File 3. Alignment used to produce ML tree in Supplementary Figure 2b

Supplementary File 4. ML tree in Supplementary Figure 2b (Newick format)

Supplementary File 5. Protein sequences containing TIR domains in Supplementary Figure 2a

Supplementary File 6. Alignment used to produce ML tree in Figure 1a

Supplementary File 7. ML tree in Figure 1a (Newick format)

Supplementary File 8. Alignment used to produce ML tree in Supplementary Figure 3

Supplementary File 9. ML tree in Supplementary Figure 3 (Newick format)

Supplementary File 10. Alignment used to produce ML tree in Supplementary Figure 4

Supplementary File 11. ML tree in Supplementary Figure 4 (Newick format)

Supplementary File 12. Alignment used to produce ML tree in Supplementary Figure 6

Supplementary File 13. ML tree in Supplementary Figure 6 (Newick format)

Supplementary File 14. Alignment used to produce ML tree in Supplementary Figure 8

Supplementary File 15. ML tree in Supplementary Figure 8 (Newick format)

Supplementary File 16. Alignment used to produce ML tree in Figure 3c

Supplementary File 17. ML tree in Figure 3c (Newick format)

Supplementary File 18. Custom Hidden Markov model based on TNP-NBARC

## Supplementary Figure Legends

**Supplementary Figure 1 TIR distribution across 39 plant species.**

**A** Total number of TIR domains predicted in plant species representing major algae and land plant taxa.

**B** Number of proteins with a TIR-NBARC-LRR (TNL) domain structure.

**C** Number of proteins with a TIR-NBARC (TN) domain structure.

**D** Number of proteins with a TIR-only architecture (<400 aa long sequences with no other predicted domains).

**Supplementary Figure 2 Complete TIR phylogeny across tested plant species.**

**A** Maximum likelihood (ML) phylogenetic tree (evolutionary model JTT+F+R10) of 2348 predicted TIR domain sequences representing major TIR families across 39 plant species (including green algae). Branches with ultrafast bootstrap (BS) support ≥95 are marked with black dots. Conserved groups with TIRs from more than one taxonomic group are highlighted with colored boxes.

**B** ML tree (evolutionary model JTT+F+R9) for 2317 predicted TIR domain sequences (same dataset as in A but excluding algal TIRs). Branches with ultrafast BS support ≥95 are marked with black dots. Conserved groups with TIRs from more than one species are highlighted with colored boxes.

**C** Same tree as in **B** with red triangles marking position of selected TIR sequences used to construct ML tree in Figure 1a.

**Supplementary Figure 3 Phylogeny of TIR-associated NBARC domains.**

ML tree (evolutionary model JTT+F+R5) for 178 NBARC domain sequences predicted as additional domains in the representative TIR protein dataset shown on the ML tree in Figure1A. Branches with ultrafast BS support ≥90 are marked with black dots. Conserved groups with TIRs from more than one species are highlighted with colored boxes.

**Supplementary Figure 4 TNP tree for aquatic plants.**

ML tree (evolutionary model PROTCATJTT, from RAxMLv8.2.9) for 201 proteins containing the NB-ARC-like domain HMM identified by HMMsearch of proteomes. Tree includes species with and without *EDS1*. Branches with BS support ≥90 are marked with black dots. The blue clade indicates TNPs.

**Supplementary Figure 5 NBARC sequence alignment and motifs.**

Amino-acid sequence alignment (from MUSCLE) of NLR proteins NB-ARC domain and TNP protein NB-ARC like domains. Black boxes highlight conserved motifs characterized due to their importance for NLR function. Above the red line are TNP protein sequences and below are TNL. CNL and RNL sequences.

**Supplementary Figure 6 TIR phylogeny including TIRs from *Chondrus crispus* and *Klebsormidium nitens*.**

ML tree (evolutionary model WAG+F+R7) for 353 predicted TIR domain sequences (same dataset as in Figure1A but including predicted TIRs from the red algae *Chondrus crispus* and the charophyte *Klebsormidium nitens*). Branches with BS support ≥90 are marked with black dots. Conserved groups with TIRs from more than one species and *Chondrus*- and *Klebsormidium*-specific groups are highlighted with colored boxes.

**Supplementary Figure 7 Alignment of predicted structures of conserved TIR-only structural comparison to TNL TIRs.**

Solved structures of the RPS4 (PDB:4c6t, chain B), RPP1 (PDB:7crc, chain C) and L6 (PDB:3ozi, chain A). TIR domains were aligned in PyMol (v3.7) to predicted structures of conserved TIR-only proteins from *Arabidopsis* (AT1G52900, AlphaFold2, UniprotID Q9C931, accessed 15 Aug 2021, alphafold.ebi.ac.uk) and rice (Os07G0566800, AlphaFold2, UniprotID Q7XIJ6, accessed 15 Aug 2021, alphafold.ebi.ac.uk). Positions of major TIR-TIR AE and DE self-association interfaces as well as the BB-loop region and catalytic glutamates are highlighted with arrows. αD helical region of conserved TIR-only proteins AT1G52900 and Os07G0566800 is likely less structured compared to RPS4, RPP1 and L6 TIRs.

**Supplementary Figure 8 EP domain phylogeny to access presence/absence of EDS1 components in plant proteomes.**

ML tree (evolutionary model JTT+F+R7) for predicted EP domain sequences. Based on phylogeny, numbers of predicted EDS1, PAD4 and SAG101 orthologues were calculated per species. Branches with BS support ≥95 are marked with black dots. Conserved groups with EP domains from EDS1, PAD4 or SAG101 are highlighted with colored boxes.

**Supplementary Figure 9 Presence-absence of TNL #1, SAG101 and orthogroups co-occurring with them across selected seed species.**

Dot plot to indicate co-occurrence of protein families. Black dots in TNP, TIR-only and TNL columns is based on phylogenetic analysis in Figure 1. For all other columns a black dot indicates presence of a protein belonging to that protein orthogroup as identified by Orthofinder, BLASTP or reciprocal tBLASTn.

**Supplementary Figure 10*TIR* gene expression in immune-triggered tissues.**

Comparison of untriggered and immune-triggered expression of *NLRs* and genes corresponding to conserved *TIR* groups in wild tobacco (*Nicotiana benthamiana*), rice (*Oryza sativa*), maize (*Zea mays*) and the liverwort *Marchantia polymorpha*. Data were taken from publicly available RNAseq experiments including immune-triggered and infected samples. Created with elements from BioRender.com.

**Supplementary Figure 11Mutant alleles of *M. polymorpha* and *N. benthamiana tnp* lines.**

**A** Representation of CRISPR/Cas9 mutant *tnp* lines in *Marchantia polymorpha*. Two sgRNA sites targeting the single *TNP* gene in *M. polymorpha* are indicated with arrows. Induced mutations are shown as alignments to the WT sequence. The two independent lines represent independent knockouts.

**B** CRISPR/Cas9 *tnp* mutant lines in *Nicotiana benthamiana*. One sgRNA site targeting each of the four *TNP* genes in *N. benthamiana* is indicated with arrows. Induced mutations are shown as alignments to the WT sequence. The two independent lines are homozygous quadruple knockouts.

## Acknowledgements

We thank Jonathan Jones (Sainsbury Laboratory, Norwich UK) for sharing *N. tabacum RNAi:EDS1* seeds. This work was supported by the Max-Planck Society (JEP, FL), Deutsche Forschungsgemeinschaft (DFG; German Research Foundation) SFB-1403–414786233 (JEP, DL, OJ, HL), DFG/Agence Nationale de la Recherche Trilateral ‘RADAR’ grant ANR-15-CE20-0016-01 (JEP, JAD), DFG SPP ‘DeCrypt’ PA-917/8-1 (JEP, CU), Germany’s Excellence Strategy CEPLAS (EXC-2048/1, Project 390686111) (JEP) the <u>Biotechnology and Biological Sciences Research Council</u> (BBSRC Doctoral Training Program BB/M011216/1 to ELB.); the <u>EC | European Research Council</u> (grant ERC-2016-STG-716233-MIREDI to KVK). OJ is a member of the International Max-Planck PhD Research School (IMPRS). We also thank Artem Pankin (formerly, MPIPZ) for advice on phylogenetic analyses and the Earlham Institute Scientific Computing group alongside the Norwich BioScience Institutes Partnership Computing infrastructure for Science (CiS) group for access to high performance computing resources.

## Author contributions

DL, OJ, ELB, KVK, and JEP conceived the project. OJ, ELB, and DL performed sequence and phylogenetic analyses. DL, OJ, KK analyzed RNAseq data. ELB predicted NLRs, OJ, JAD, CU, KM, HN developed CRISPR/Cas9 mutant lines. OJ, HLL, DL, FL, ELB performed immunology assays. OJ, ELB, DL, KVK and JEP analysed the data. OJ, DL and JEP wrote the manuscript with contributions from ELB and KVK. All authors commented on the manuscript.

