## Supplementary material for "Differential *EDS1* requirement for cell death activities of plant TIR-domain proteins": Supp_Figures_v13.pdf

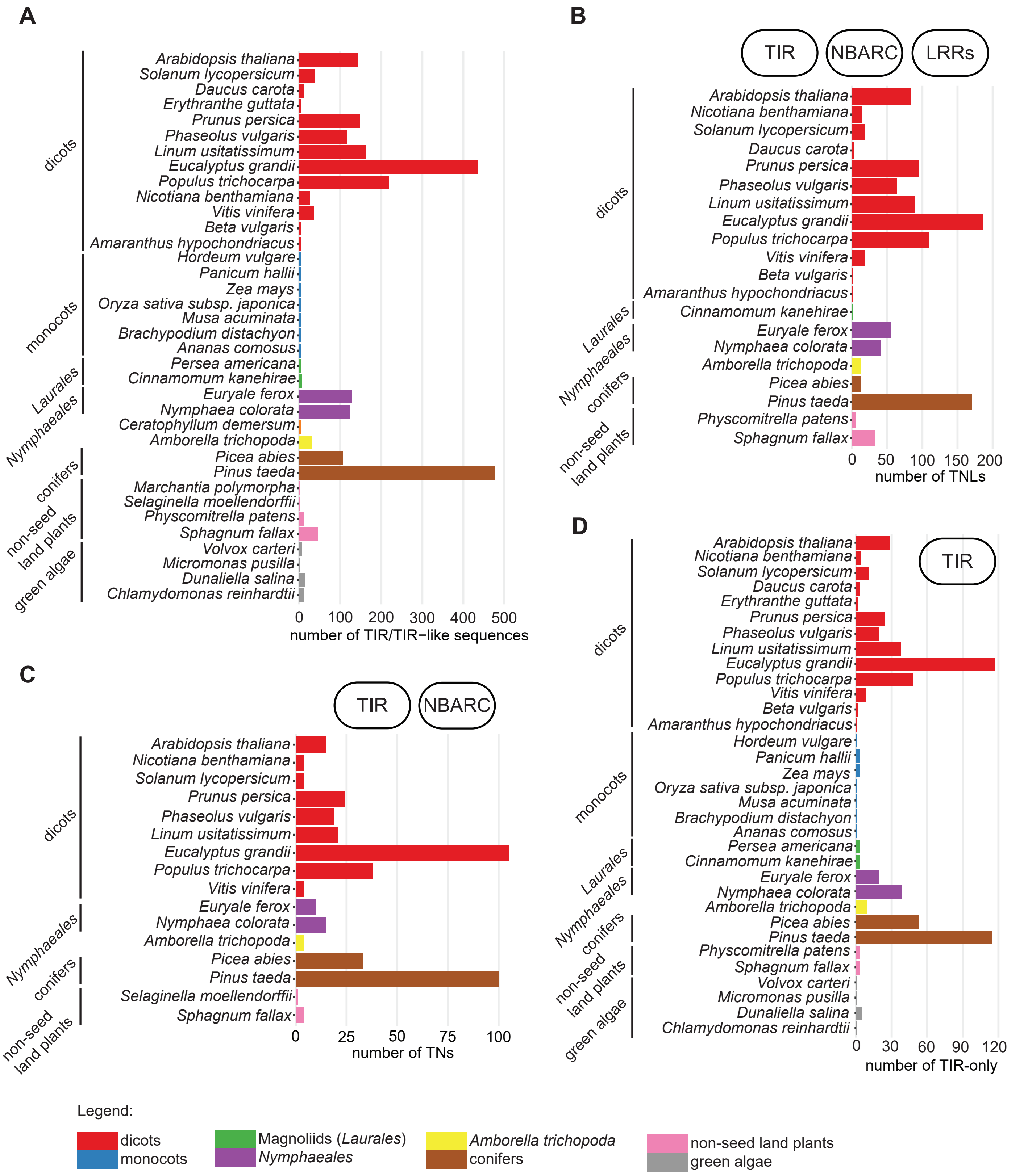


**Supplementary Figure 1. TIR distribution across 39 plant species.**

**A** Total number of TIR domains predicted in plant species representing major algae and land plant taxa.

**B** Number of proteins with a TIR-NBARC-LRR (TNL) domain structure.

**C** Number of proteins with a TIR-NBARC (TN) domain structure.

**D** Number of proteins with a TIR-only architecture (<400 aa long sequences with no other predicted domains).

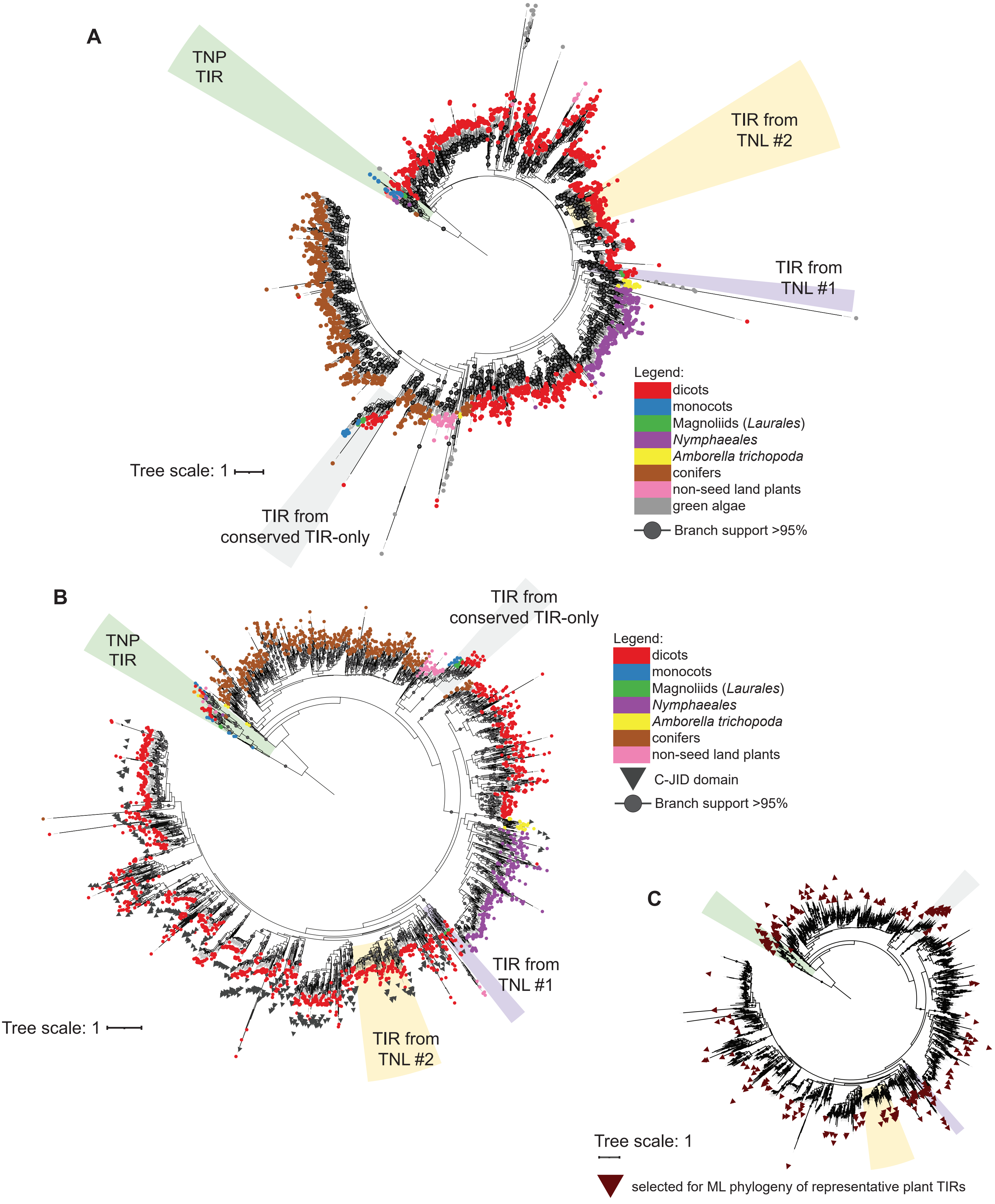
A
TNP TIR
TIR from TNL #2
TIR from TNL #1
TIR from conserved TIR-only
Tree scale: 1
Legend:
dicots
monocots
Magnoliids (Laurales)
Nymphaeales
Amborella trichopoda
conifers
non-seed land plants
green algae
Branch support >95%
B
TNP TIR
TIR from conserved TIR-only
Legend:
dicots
monocots
Magnoliids (Laurales)
Nymphaeales
Amborella trichopoda
conifers
non-seed land plants
C-JID domain
Branch support >95%
TIR from TNL #1
TIR from TNL #2
Tree scale: 1
C
Tree scale: 1
selected for ML phylogeny of representative plant TIRs

**Supplementary Figure 2. Complete TIR phylogeny across tested plant species.**

**A** ML tree (from IQ-TREE, evolutionary model JTT+F+R10) of 2348 predicted TIR domain sequences representing major TIR families across 39 plant species (including green algae). Branches with BS support ≥95 are marked with black dots. Conserved groups with TIRs from more than one species are highlighted with colored boxes.

**B** ML tree (from IQ-TREE, evolutionary model JTT+F+R9) for 2317 predicted TIR domain sequences (same dataset as in A but excluding algal TIRs). Branches with BS support ≥95 are marked with black dots. Conserved groups with TIRs from more than one species are highlighted with colored boxes.

**C** Same tree as in B with red triangles marking position of selected TIR sequences used to construct ML tree in Figure 1a.

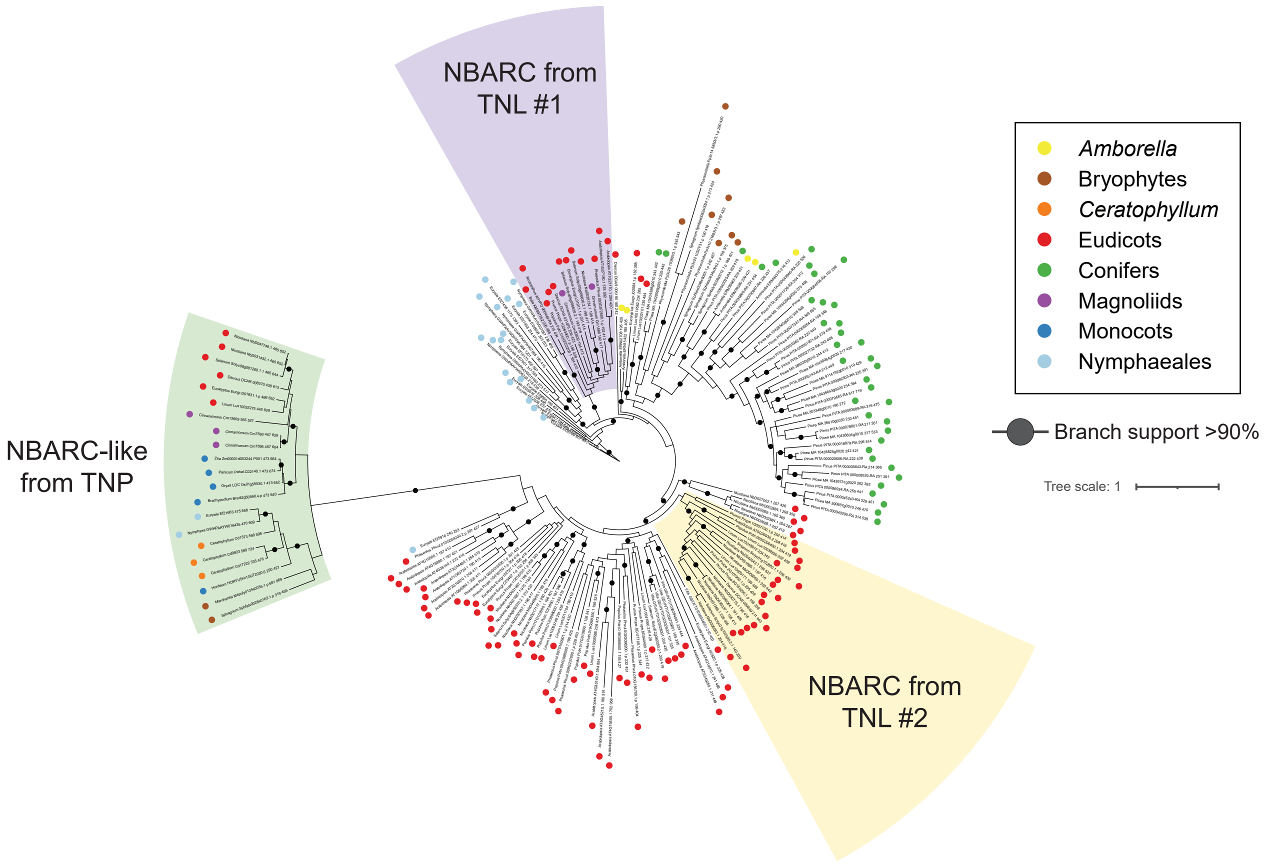


**Supplementary Figure 3 Phylogeny of TIR-associated NBARC domains.**
ML tree (from IQ-TREE, evolutionary model JTT+F+R5) for 178 NBARC domain sequences predicted as additional domains in the representative TIR protein dataset shown on the ML tree in Figure1A. Branches with BS support ≥90 are marked with black dots. Conserved groups with TIRs from more than one species are highlighted with colored boxes.

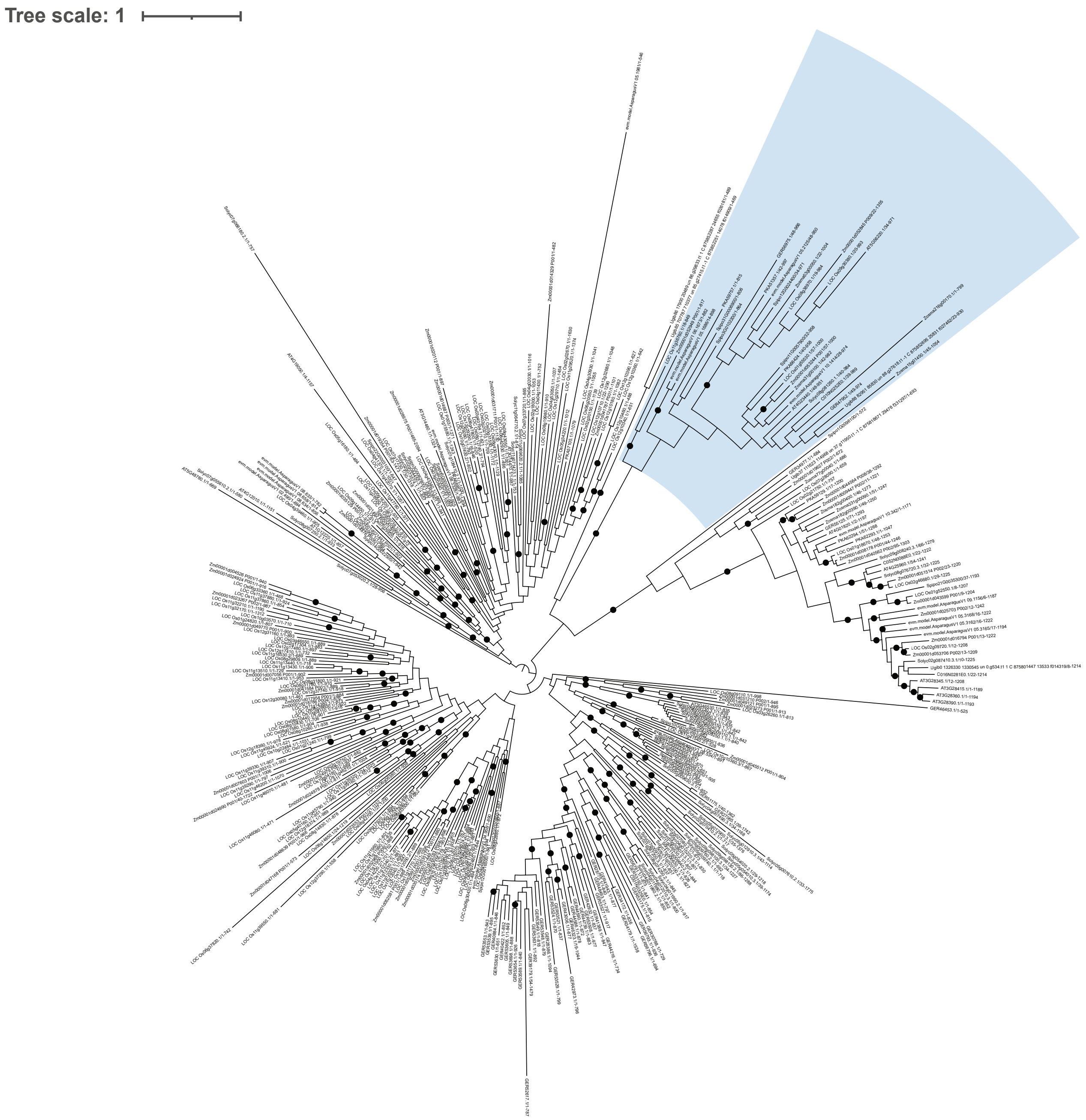


**Supplementary Figure 4. TNP tree for plants including species without EDS1.**
ML tree (from RAxMLv8.2.9, evolutionary model PROTCATJTT) for 201 proteins containing the NB-ARC-like domain identified by HMMsearch of proteomes, tree includes species with and without EDS1. Branches with BS support ≥90 are marked with black dots. The blue clade indicates TNP containing group.

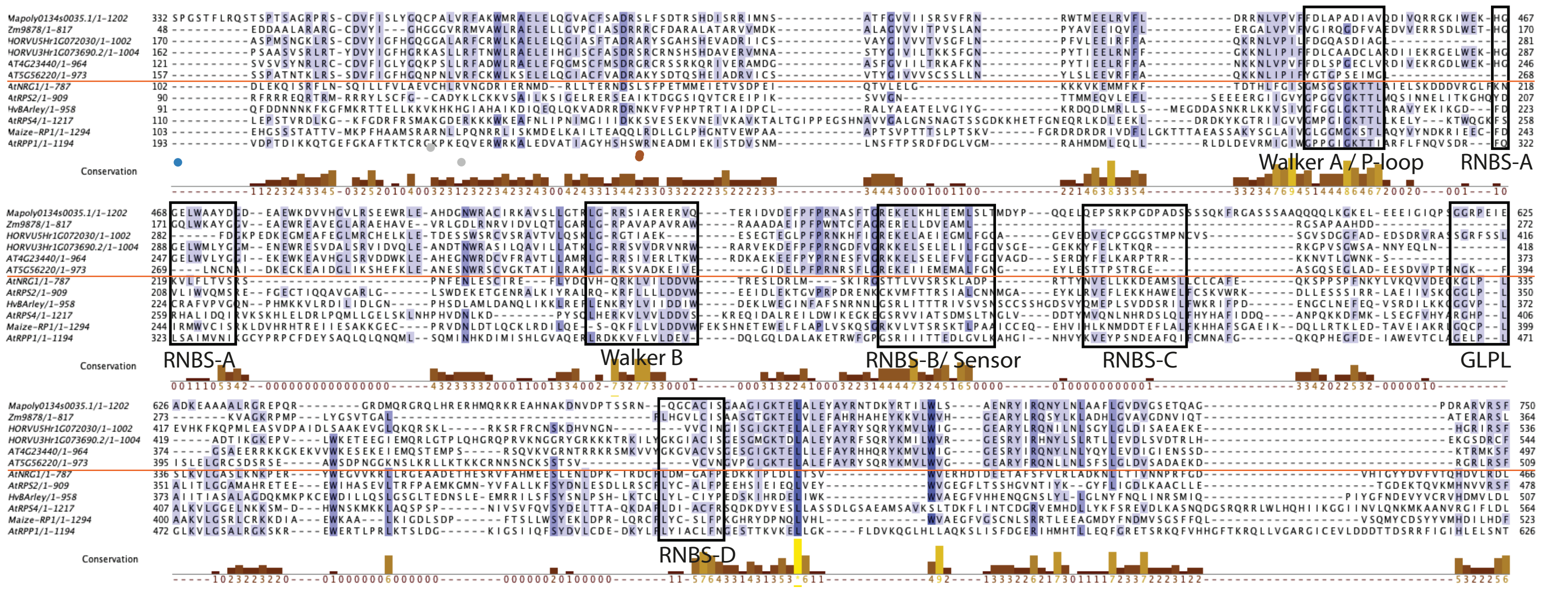


**Supplementary Figure 5. NBARC sequence alignment and motifs.**
Amino-acid sequence alignment (from MUSCLE) of NLR proteins NB-ARC domain and TNP protein NBARC like domains. Black boxes highlight conserved motifs characterized due to their importance for NLR function. Above the red line are TNP protein sequences and below are TNL. CNL and RNL sequences.

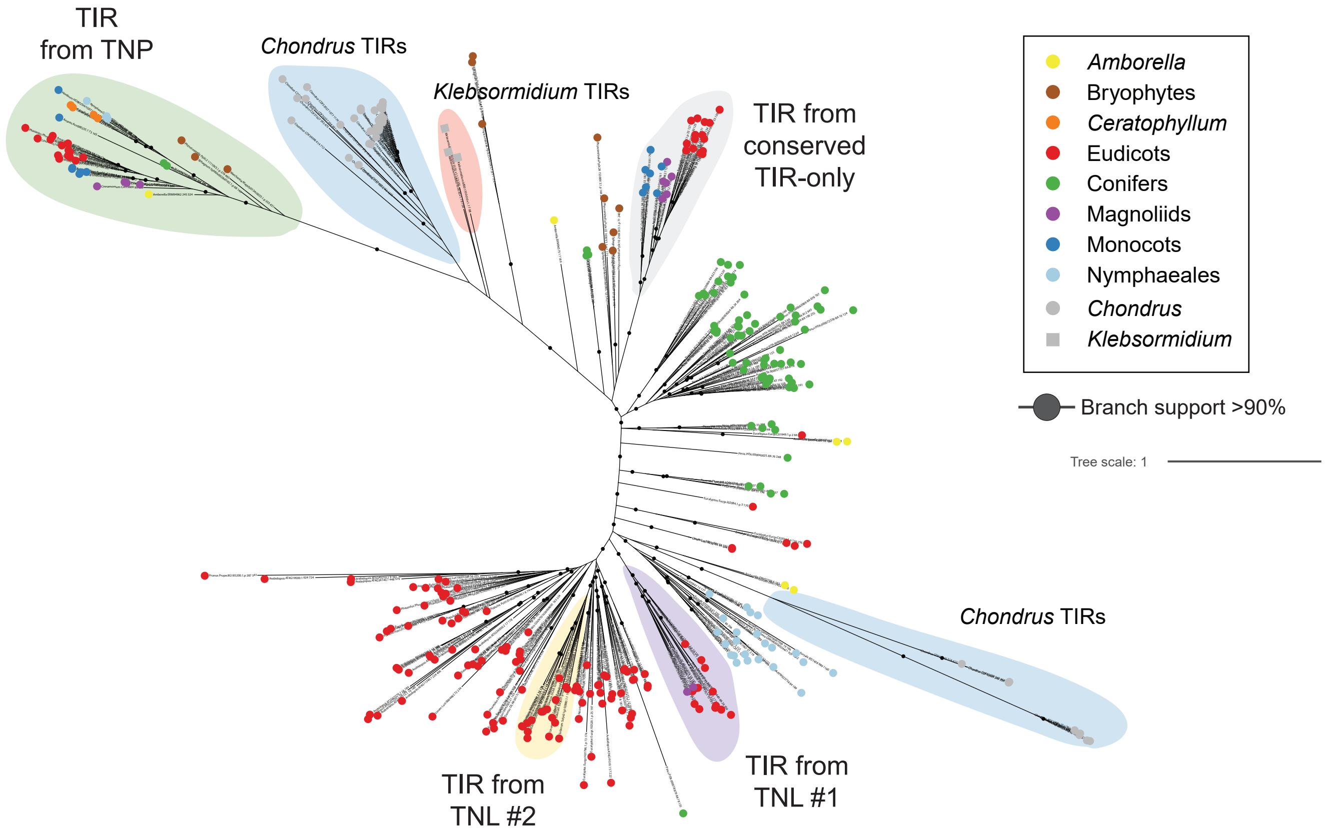


**Supplementary Figure 6. TIR phylogeny including TIRs from *Chondrus* and *Klebsormidium*.** ML tree (from IQ-TREE, evolutionary model WAG+F+R7) for 353 predicted TIR domain sequences (same dataset as in Figure1A but including predicted TIRs from the red algae *Chondrus crispus* and the charophyte *Klebsormidium nitens*). Branches with BS support ≥90 are marked with black dots. Conserved groups with TIRs from more than one species and *Chondrus*- and *Klebsormidium*-specific groups are highlighted with colored boxes.

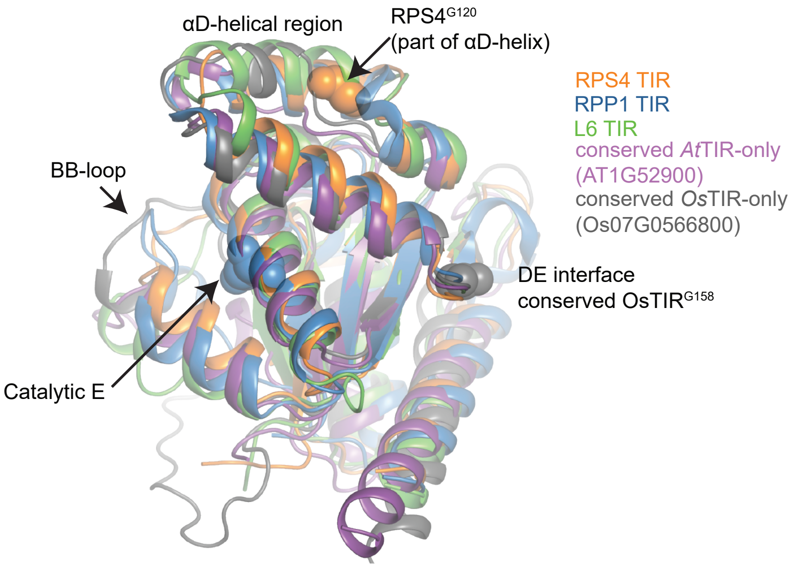


**Supplementary Figure 7. Alignment of predicted structures of conserved TIR-only structural comparison to TNL TIRs.**

Solved structures of the RPS4 (PDB:4c6t, chain B), RPP1 (PDB:7crc, chain C) and L6 (PDB:3ozi, chain A). TIR domains were aligned in PyMol (v3.7) to predicted structures of conserved TIR-only proteins from *Arabidopsis* (AT1G52900, AlphaFold2, UniprotID Q9C931, accessed 15 Aug 2021, alphafold.ebi.ac.uk) and rice (Os07G0566800, AlphaFold2, UniprotID Q7XIJ6, accessed 15 Aug 2021, alphafold.ebi.ac.uk). Positions of major TIR-TIR AE and DE self-association interfaces as well as the BB-loop region and catalytic glutamates are highlighted with arrows. αD helical region of conserved TIR-only proteins AT1G52900 and Os07G0566800 is likely less structured compared to RPS4, RPP1 and L6 TIRs.

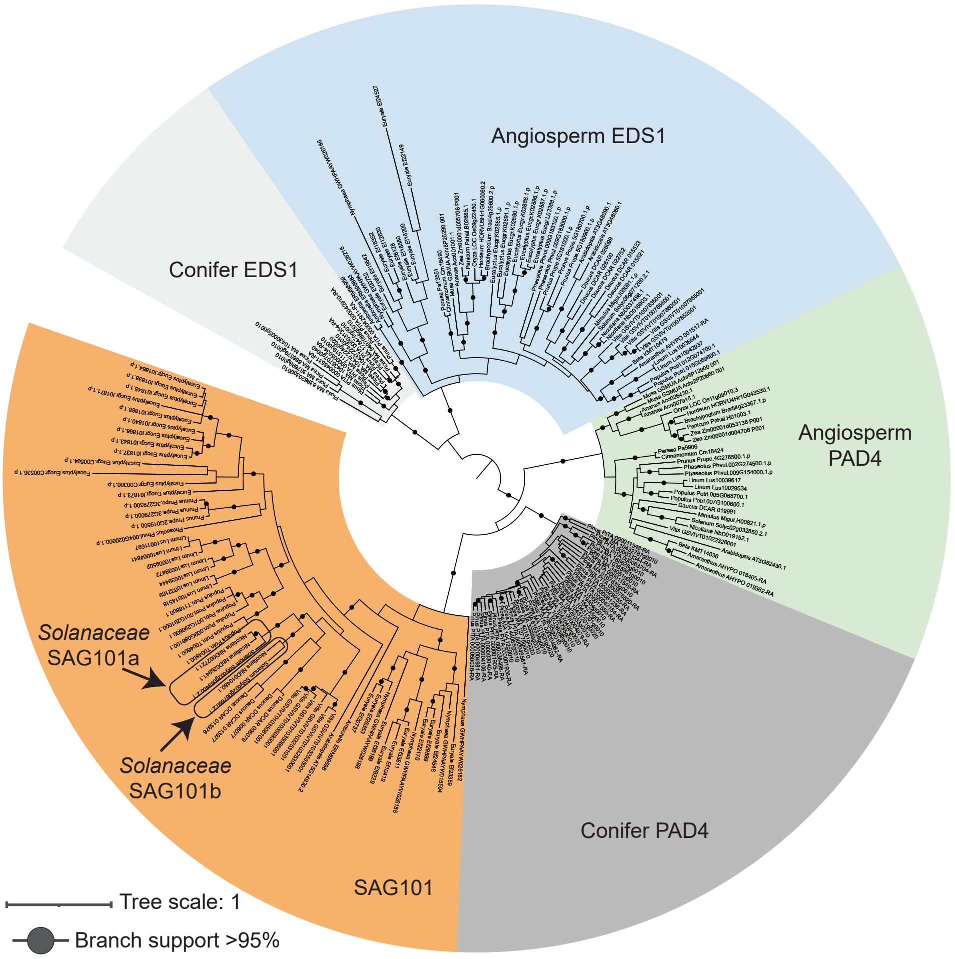


**Supplementary Figure 8. EP domain phylogeny to access presence/absence of EDS1 components in plant proteomes.**
ML tree (from IQ-TREE, evolutionary model JTT+F+R7) for predicted EP domain sequences. Based on phylogeny, numbers of predicted EDS1, PAD4 and SAG101 orthologues were calculated per species. Branches with BS support ≥95 are marked with black dots. Conserved groups with EP domains from EDS1, PAD4 or SAG101 are highlighted with colored boxes.

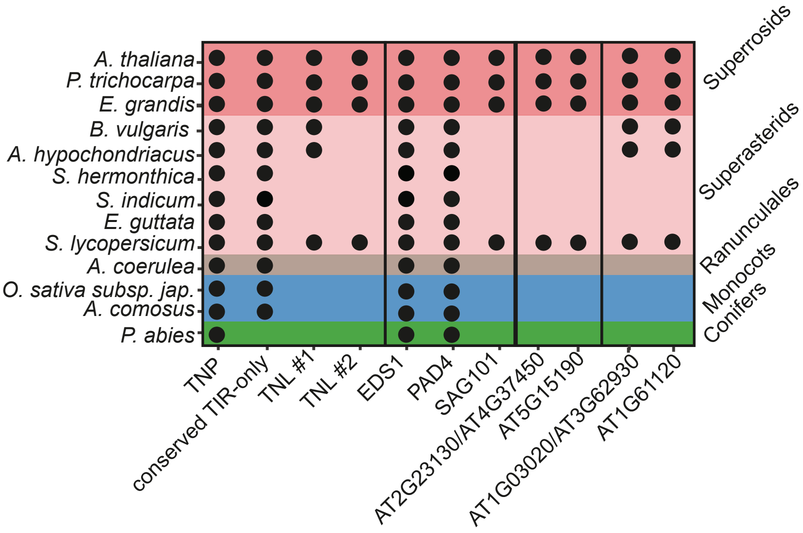


**Supplementary Figure 9. Presence-absence of TNL #1, SAG101 and orthogroups co-occurring with them across selected seed plant species.**

Dot plot to indicate co-occurrence of protein families. Black dots in TNP, TIR-only and TNL columns is based on phylogenetic analysis in Figure 1. For all other columns a black dot indicates presence of a protein belonging to that protein orthogroup as identified by Orthofinder, BLASTP or reciprocal tBLASTn.

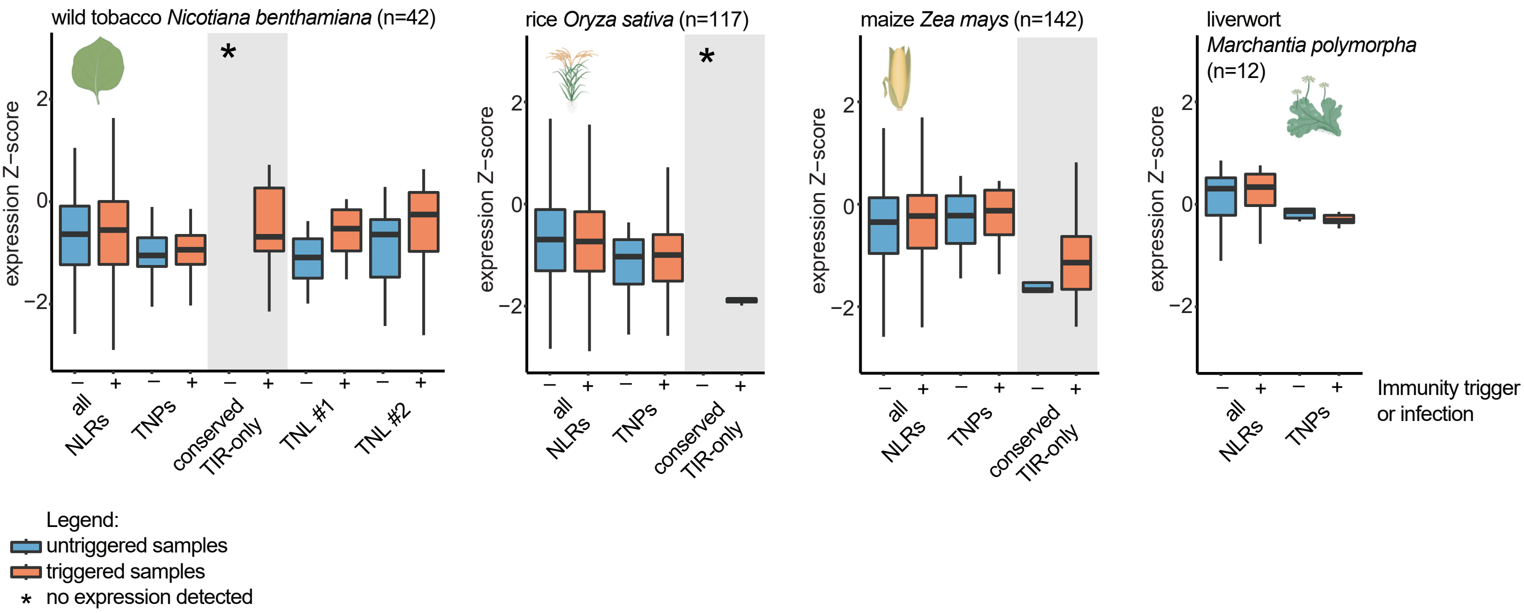


**Supplementary Figure 10. *TIR* gene expression in immune-triggered tissues.**
Comparison of untriggered and immune-triggered expression of NLRs and genes corresponding to conserved TIR groups in wild tobacco (*Nicotiana benthamiana*), rice (*Oryza sativa*), maize (*Zea mays*) and the liverwort *Marchantia polymorpha*. Data were taken from publicly available RNAseq experiments (Table S4) including immune-triggered and infected samples. Created with elements from BioRender.com.

**A**

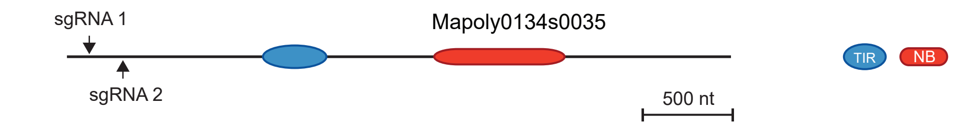


```
Mapoly0134s0035    sgRNA 1
WT:     GCTCTCTCGTCGTCCTCGTCGTCGTCGCTCGCGTGCCC
tnp1:   GCTCTCTCGTCGTCCTCGT-GTCGTCGCTCGCGTGCCC
tnp2:   GCTCTCTCGTCGTCCTCGTCGTCGTCGCTCGCGTGCCC

Mapoly0134s0035    sgRNA 2
WT:     CGCCCTCGTCGCCCTCCG---------TCTCTCACAGCATTGC
tnp1:   CGCCCTCGTCGCCCTCCG---------TCTCTCACAGCATTGC
tnp2:   CGCCCTCGTCGCCCTCCTCGCGTCGTCTCTCTCACAGCATTGC
```

**B**

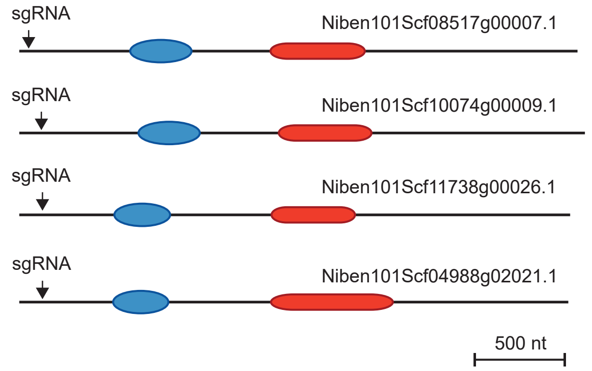


```
Niben101Scf08517g00007.1
WT:      CCACTGTTCTTCCTTCAGG-TCAGCAGAAATTGGTAGC
tnp-q1:  CCACTGTTCTTCCTTCAGGGTCAGCAGAAATTGGTAGC
tnp-q2:  CCACTGTTCTTCCT------TCAGCAGAAATTGGTAGC

Niben101Scf10074g00009.1
WT:      CCACAGTTCTTCCTTCA-GGTCAGCAGAAATTGGTAGC
tnp-q1:  CCACAGTTCTTCCTTCATGGTCAGCAGAAATTGGTAGC
tnp-q2:  CCACAGTTCTTCCTTCA--GTCAGCAGAAATTGGTAGC

Niben101Scf11738g00026.1
WT:      TCTCCCAGGTCACCCAAGT-CCCTGATATCAACATATTCA
tnp-q1:  TCTCCCAGGTCACCCAAGTTCCCTGATATCAACATATTCA
tnp-q2:  TCTCCCAGGTCACCCAAGT-- (71 bp deletion) --

Niben101Scf04988g02021.1
WT:      CCAGGTCACCCAAG--------------------TCCCTGATATCAACATAT
tnp-q1:  CCAGGTCACCCAAG-(231 bp insertion)-TCCCTGATATCAACATAT
tnp-q2:  CCAGGTCACCCAAGC-------------------TCCCTGATATCAACATAT
```

**Supplementary Figure 11. Mutant alleles of *M. polymorpha* and *N. benthamiana tnp* lines.**

**A** Representation of CRISPR/Cas9 mutant tnp lines in *Marchantia polymorpha*. Two sgRNA sites targeting the single *TNP* gene in *M. polymorpha* are indicated with arrows. Induced mutations are shown as alignments to the WT sequence. The two independent lines represent independent knockouts.

**B** CRISPR/Cas9 tnp mutant lines in *Nicotiana benthamiana*. One sgRNA site targeting each of the four *TNP* genes in *N. benthamiana* is indicated with arrows. Induced mutations are shown as alignments to the WT sequence. The two independent lines are homozygous quadruple knockouts.
